# Vitamin K-dependent carboxylation in osteoblasts regulates bone resorption through GAS6 in male mice

**DOI:** 10.1101/2025.09.08.673085

**Authors:** Monica Pata, Diep Ngoc Thi Pham, Julie Lacombe, B. Ashok Reddy, Young Woong Kim, Abeer Gamal Ali Ahmed, Monzur Murshed, Mathieu Ferron

**Author notes:** Correspondence to: Mathieu Ferron, PhD, Institut de Recherches Cliniques de Montréal 110 Ave. des Pins O., Montréal, QC, H2W 1R7, Canada. Author’s. **Contributions** M.F. and M.P. assembled the figures. M.F. wrote the manuscript with suggestions from the other authors. M.P., M.F., D.N.T.P., J.L., B.A.R. and Y.W.K., collected and analyzed data. M.F. conceived and designed the project and provided supervision to M.P., D.N.T.P., J.L., B.A.R. and Y.W.K. A.G.A.A. and M.M. collected and analyzed the mCT data. M.F. is the guarantor of this work and, as such, had full access to all the data in the study and takes responsibility for the integrity of data and the accuracy of data analysis.

## Abstract

Studies in humans suggest that vitamin K is involved in the regulation of bone remodeling, but the precise mechanism at play remains unknown. In cells, vitamin K functions as a co-factor for the γ-glutamyl carboxylase (GGCX), an enzyme responsible for the conversion of glutamic acid residues (Glu) into γ-carboxyglutamic acid (Gla) residues in secreted proteins. We aim here at determining the role of γ-carboxylation in bone remodeling and at identifying the Gla protein(s) involved. We show that male mice lacking γ-carboxylation specifically in osteoblasts (*Ggcx^flox/flox^*;*OCN-Cre*) have increased bone mass at 6 months of age due to a reduced number of multinucleated bone resorbing osteoclasts. In co-culture experiments, *Ggcx*-deficient osteoblasts were less effective than control osteoblasts at supporting osteoclast formation. Among known Gla proteins, we identify GAS6 as an osteoblast-secreted γ-carboxylated factor which signals to differentiating osteoclasts. The GAS6 receptors MerTK and AXL are expressed in pre-osteoclasts and pharmacological inhibitors of AXL and MerTK block osteoclast generation in co-culture. Conversely, recombinant γ-carboxylated GAS6 dose-dependently increases the size of osteoclasts and the number of nuclei per osteoclast in culture. GAS6 marginally affected the induction of osteoclast-specific genes during osteoclast differentiation but significantly increased pre-osteoclast fusion. Finally, increasing bone marrow GAS6 level in transgenic male mice was sufficient to increase the number and size of osteoclasts and to decrease bone mass. This work identifies GAS6 as a novel osteoblast-derived vitamin K-dependent protein regulating osteoclast maturation.

## Introduction

The acquisition and maintenance of an adequate skeleton depend on the process of bone turnover, which requires the activity of two cell types: the osteoclast, responsible for the resorption of the mineralized bone extracellular matrix (ECM) and the osteoblast, responsible for the synthesis, secretion and mineralization of the bone ECM. An imbalance between the activities of these two cell types can result in severe bone diseases such as osteoporosis. Like many chronic aging disorders, osteoporosis appears to be a multifactorial disease. Hence, while genetic variants may contribute to the physiopathology of osteoporosis ^1,2^, epigenetic, environmental and behavioral factors have also been implicated ^3,4^. For instance, several micronutrients including vitamins are known to play a role in the acquisition and the maintenance of a healthy skeleton and could therefore influence the etiology of osteoporosis ^4,5^. This is evidenced by the profound impact that vitamins C, D or B12 deficiencies has on bone density or quality in humans ^6–9^. Although isolated vitamin K (VK) deficiency in adult is a rare condition, longitudinal prospective studies have reported that lower VK intake or serum levels were associated with decreased bone mineral density (BMD) ^10,11^ or increased fracture risk in postmenopausal women ^12–14^.

There are two naturally occurring forms of VK: VK_1_ or phylloquinone, present in green vegetable, and VK_2_ or menaquinones (e.g., MK-4 and MK-7), present in animals and fermented foods ^15^. VK_1_ is an essential micronutrient which can only be obtained through alimentation, but it can be converted to MK-4 in mammalian cells by the enzyme UbiA prenyltransferase domain containing 1 (UBIAD1). Bone contains both VK_1_ and MK-4 ^16,17^, and a number of clinical trials have tested the effects of VK_1_ or VK_2_ supplementation on BMD or fracture risk, which led to conflicting results ^15^. Although some studies reported that MK-4, MK-7, or VK supplementation increased femoral or vertebral bone mineral density (BMD) and/or reduced fracture risk, others found no significant effect ^18–22^. A few meta-analyses of interventional studies suggest that VK supplementation had a significant and positive effect on BMD at lumbar spine and forearm ^18,19,23^, while another indicated that VK prevents fractures without impacting BMD^24^. Therefore, whether VK is directly implicated in the regulation of bone mass or quality remains controversial. Moreover, the molecular and cellular mechanisms through which VK influences bone density and/or quality remain uncharacterized.

In vertebrates, the only well described cellular function of VK is to serve as a co-factor during the γ-carboxylation reaction that converts glutamic acid (Glu) residues to γ-carboxyglutamic acid (Gla) residues in specific proteins in the endoplasmic reticulum ^25^. Gamma-carboxylated proteins (also called “Gla” proteins) are either secreted or transmembrane polypeptides with the potential of having endocrine, paracrine or autocrine functions. Two enzymes are involved in γ-carboxylation: γ-glutamyl carboxylase (γ-carboxylase or GGCX) and vitamin K oxidoreductase (VKORC1) ^26^. γ-carboxylase requires reduced VK (VKH_2_) as an essential cofactor, which upon carboxylation is oxidized to VK epoxide (VKO). VKO is next reconverted to VKH_2_ by VKORC1.

Osteocalcin (Ocn) is an osteoblast-derived Gla protein present at high concentration in the bone ECM. Because γ-carboxylated Ocn has great affinity for hydroxyapatite, the mineral component of bone ECM, it was originally believed to play a role in bone mineralization ^27^. The analysis of *Ocn*-deficient mice or rats (*Ocn^-/-^*) suggested that the absence of osteocalcin may slightly increase bone formation and bone density ^28,29^. However, two other studies involving different strains of *Ocn^-/-^* mice did not detect any impact on bone homeostasis ^30,31^. Altogether, these genetic data do not support a major role for Ocn in bone mass accrual or turnover and suggest that the mechanism and the Gla protein(s) through which VK may influence bone health remain to be characterized.

We aim here at addressing specifically the role of γ-carboxylation in bone remodeling in vivo. We show that the osteoblast-specific inactivation of *Ggcx* in mice is associated with an increased bone density resulting from a reduced number of osteoclasts without significant impact on bone formation. We establish that *Ggcx*-deficient osteoblasts are less effective at supporting osteoclast formation ex vivo and identify GAS6 as an osteoblast-secreted γ-carboxylated factor which signals to osteoclast precursors. Finally, using cell-based assays and a gain-of-function model in mice, we show that γ-carboxylated GAS6 promotes pre-osteoclast fusion. These data reveal an unexpected role for γ-carboxylation in coupling bone resorption to bone formation and provides new insights on the function of VK in bone homeostasis.

## Results

### The enzymes of the VK cycle are predominantly expressed in osteoblasts

To identify the bone cell type(s) mediating the potential impact of VK on bone, we examined the expression of GGCX and VKORC1 in osteoblasts and osteoclasts at the mRNA and protein levels. Quantitative PCR (qPCR) revealed that *Ggcx* and *Vkorc1* expression levels are about ten-to forty-fold higher in mouse proliferating non-mineralized osteoblasts (pre-OB) and in mineralized osteoblasts (OB) compared to bone marrow derived monocytes (BMMC) or fully differentiated osteoclasts (OCL) (**Fig.1A**). Western blotting confirmed that GGCX and VKORC1 are more abundant in primary osteoblast cultures compared to osteoclasts. Interestingly, the expression level of GGCX and VKORC1 in osteoblasts was comparable to the one observed in liver where these enzymes are known to play a critical role in the γ-carboxylation and activation of several coagulation factors (**Fig. 1B**).

**Figure 1.**
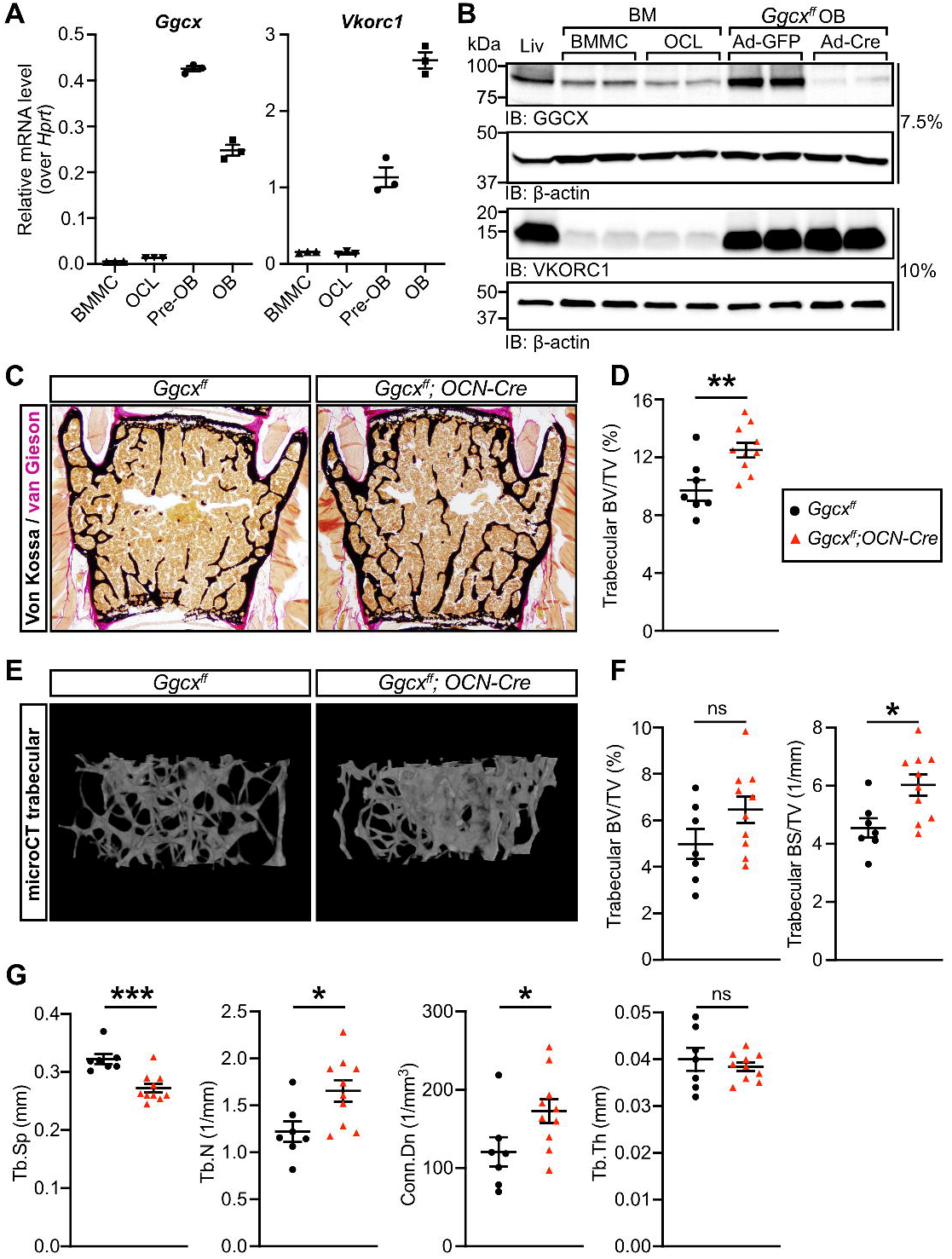
Increased bone mass in mice lacking. γ**-carboxylation in osteoblasts. (A)** Gene expression analysis by qPCR of *Ggcx* and *Vkorc1* in bone marrow derived monocytes (BMMC), osteoclasts (OCL), proliferating pre-osteoblasts (pre-OB) and mineralized osteoblast (OB) cultures (n=3). **(B)** Protein expression in liver (Liv) and bone cells by Western blot. GGCX was analyzed on a 7.5% SDS Tris Glycine gel using 20μg of extracts, while VKORC1 was resolved on a 10% SDS Tris Tricine gel using 10μg of extracts. **(C-G)** Six-month-old *Ggcx^ff^* and *Ggcx^ff^;OCN-Cre* male mice were analyzed (n=7-10). **(C)** Representative pictures of sections from lumbar vertebrae stained with von Kossa and van Gieson. **(D)** Quantification of trabecular bone volume over tissue volume (BV/TV) from the L4 and L5 lumbar vertebrae sections. **(E)** Representative μCT images of the distal femur trabecular bones. **(F)** Quantification of trabecular bone volume (BV/TV) and trabecular bone surface density (BS/TV) from the μCT data. **(G)** Trabecular bone µCT derived data. Tb.Sp., Tb.N., and Tb.Th., trabecular spacing, number, and thickness respectively; Conn.Dn., connectivity density. Unpaired, 2-tailed Student’s t test was used in (D), (F), and (G). ***p < 0.001, **p < 0.01, *p < 0.05, ns: non-significant.

### Inactivation of GGCX in osteoblasts results in increased bone density in mice

Given the predominant expression of the VK-cycle enzymes in osteoblasts, we inactivated *Ggcx* specifically in this cell type and investigated the impact on bone remodeling. For this purpose, *Ggcx^fl/fl^* mice were bred with *Osteocalcin-Cre* (OCN-Cre) mice expressing the Cre recombinase in mature osteoblasts only ^32^. We previously reported that *Ggcx* was specifically and efficiently inactivated in osteoblasts in *Ggcx^fl/fl^*;*OCN-Cre* mice ^33^. The percentage of circulating carboxylated osteocalcin, an osteoblast-specific VK-dependent protein, was reduced by 80% to 90% between 2 and 6 months of age in these mice, confirming efficient inactivation of VK-dependent carboxylation in osteoblasts in vivo (**Supplementary Fig. 1**). In a previous study, we reported that 3-month-old *Ggcx^fl/fl^*;*OCN-Cre* male mice had normal trabecular bone density ^33^. Since bone mass decreases with aging, we characterized the bone parameters of the *Ggcx^fl/fl^*;*OCN-Cre* male mice at 6 months of age when bone mass is declining in this species. Histological analysis of lumbar vertebrae indicated that trabecular bone density was significantly increased by about 30% in 6-month-old *Ggcx^fl/fl^*;*OCN-Cre* males (**Fig. 1C-D**). Microcomputed tomography (µCT) analysis indicated a significant increase in trabecular bone surface density (BS/TV), in the femur of *Ggcx^fl/fl^*;*OCN-Cre* mice, although trabecular bone density (BV/TV) was not significantly changed, likely due to higher variation (**Fig. 1E-F**). In addition, trabecular spacing (Tb.Sp) was significantly reduced, while trabecular number (Tb.N) and connectivity density (Conn.Dn), a computational measure of the inter-connectivity among trabeculae, were both significantly increased (**Fig. 1G**). Trabecular thickness (Tb. Th) was unaffected in *Ggcx^fl/fl^*;*OCN-Cre* femurs (**Fig. 1G**). Altogether, these data indicate that the absence of VK-dependent carboxylation in osteoblasts is associated with an increased bone mass at 6 months of age in male mice.

### Decreased bone resorption in *Ggcx^fl/fl^*;*OCN-Cre* mice

To get insight into the cellular mechanism behind the increased bone mass phenotype of the *Ggcx^fl/fl^*;*OCN-Cre* mice, static and dynamic bone histomorphometry analyses were performed at 6 months of age. Mineral apposition rate (MAR), measured using double calcein labeling, was slightly but significantly reduced in absence of GGCX in osteoblasts (**Fig.2A-B**). However, bone formation rate (BFR/BS), osteoblast number (N.Ob/B.Pm) and osteoblast surface (Ob.S/BS) were not significantly affected (**Fig.2C-E**). In contrast, both osteoclast number (N.Oc/B.Pm) and surface (Oc.S/BS) were significantly decreased in *Ggcx^fl/fl^*;*OCN-Cre* mice (**Fig. 2F-H**). Decrease circulating carboxy-terminal collagen crosslinks (CTX) independently confirmed reduced osteoclast activity in the same animals (**Fig.2I**). Overall, these analyses suggest that the increased bone mass observed in the absence of γ-carboxylation in osteoblasts is mainly driven by a reduction in osteoclastic bone resorption.

**Figure 2.**
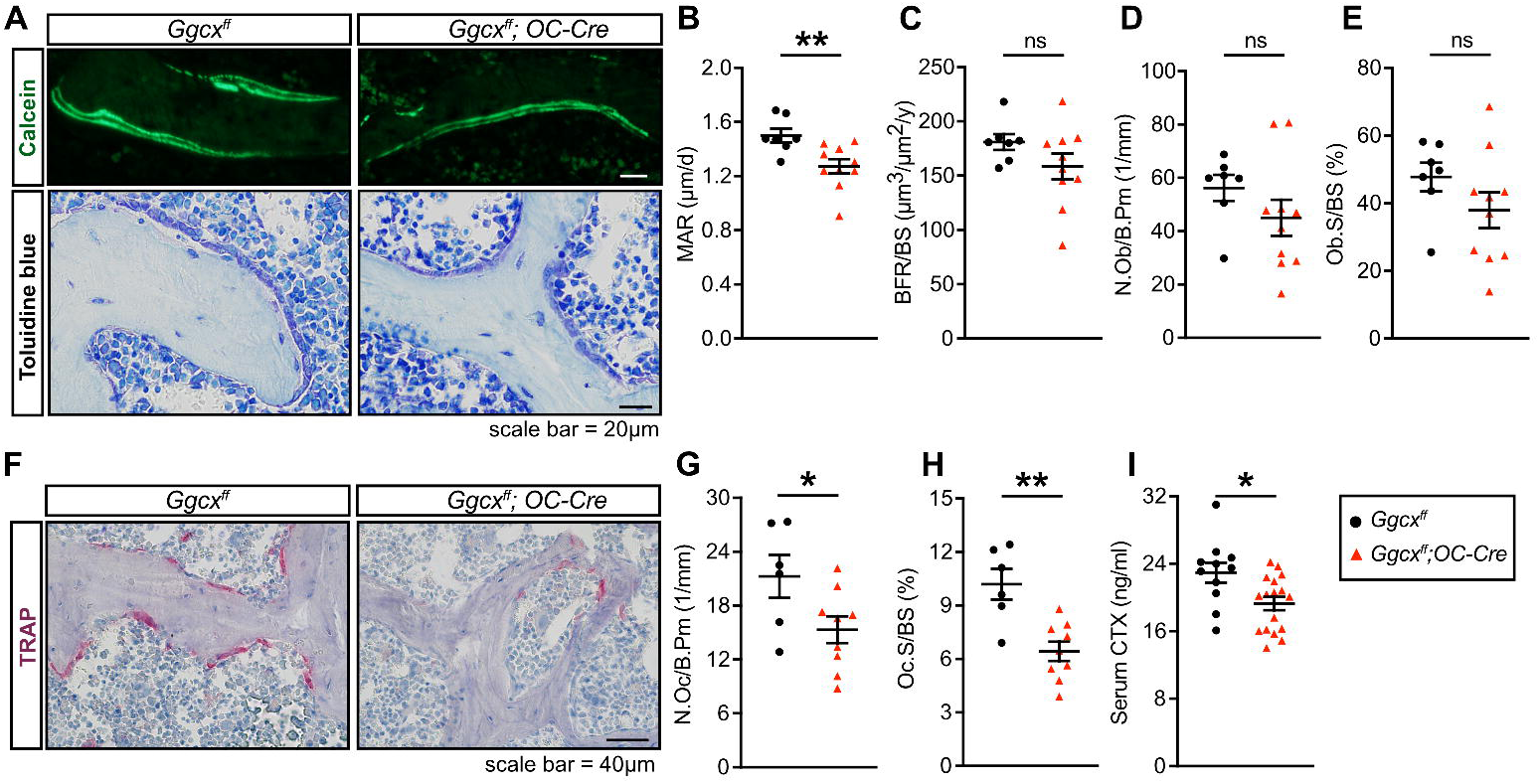
Reduced osteoclast number and surface in *Ggcx^ff^;OCN-Cre* mice. (A-H) Bone histomorphometry analysis of lumbar vertebrae in six-month-old *Ggcx^ff^* and *Ggcx^ff^;OCN-Cre* male mice (n=6-10). **(A)** Representative pictures of calcein double labeling and toluidine blue staining. **(B)** Mineral apposition rate (MAR). **(C)** Bone formation rate over bone surface (BFR/BS). **(D)** Number of osteoblasts per bone perimeter (N.Ob/B.Pm). **(E)** Osteoblast surface over bone surface (Ob.S/BS). **(F)** Representative pictures of TRAP staining. **(G)** Number of osteoclasts per bone perimeter (N.Oc/B.Pm). **(H)** Osteoclast surface over bone surface (Oc.S/BS). **(I)** Fasting serum CTx levels (n=12-17). Results represent the mean ± SEM. Unpaired, 2-tailed Student’s t test was used in (B-E) and (G-I). **p < 0.01, *p < 0.05, ns: non-significant.

### Decreased osteoclastogenesis in absence of *Ggcx* in osteoblasts ex vivo

Osteoclast differentiation is dependent on factors such as RANKL and M-CSF, produced by cells of the osteoblast lineage including pre-osteoblasts and osteocytes. Because of the reduction in osteoclast number in the *Ggcx^fl/fl^*;*OCN-Cre* mice, we hypothesized that *Ggcx*-deficient osteoblasts may be less efficient at supporting osteoclast differentiation or maturation. This was tested using co-cultures of wildtype bone marrow cells with either osteoblasts lacking GGCX (i.e., *Ggcx^fl/fl^* osteoblasts transduced with a CRE expressing adenovirus) or control osteoblasts (i.e., *Ggcx^fl/fl^* osteoblasts transduced with a GFP expressing adenovirus). We previously reported very efficient inactivation of GGCX in osteoblasts at the mRNA and protein levels using this approach ^33^ (see also **Fig. 1B**). The coculture was performed in the presence of 1,25 vitamin D_3_ (VitD3) and prostaglandin E_2_ (PGE2), which promote the expression of pro-osteoclastogenic factors by osteoblasts. These experiments indicated that the number of osteoclasts was significantly reduced by approximately threefold in the co-cultures involving GGCX-deficient osteoblasts (**Fig.3A-B**). This result suggests that γ-carboxylation in osteoblasts may regulate the production or the activity of a pro-osteoclastogenic factor. Importantly, the absence of osteocalcin (OCN), a γ-carboxylated protein specifically secreted by osteoblasts and osteocytes, did not affect the number of osteoclasts obtained from WT bone marrow cells (**Fig. 3C-D**).

**Figure 3.**
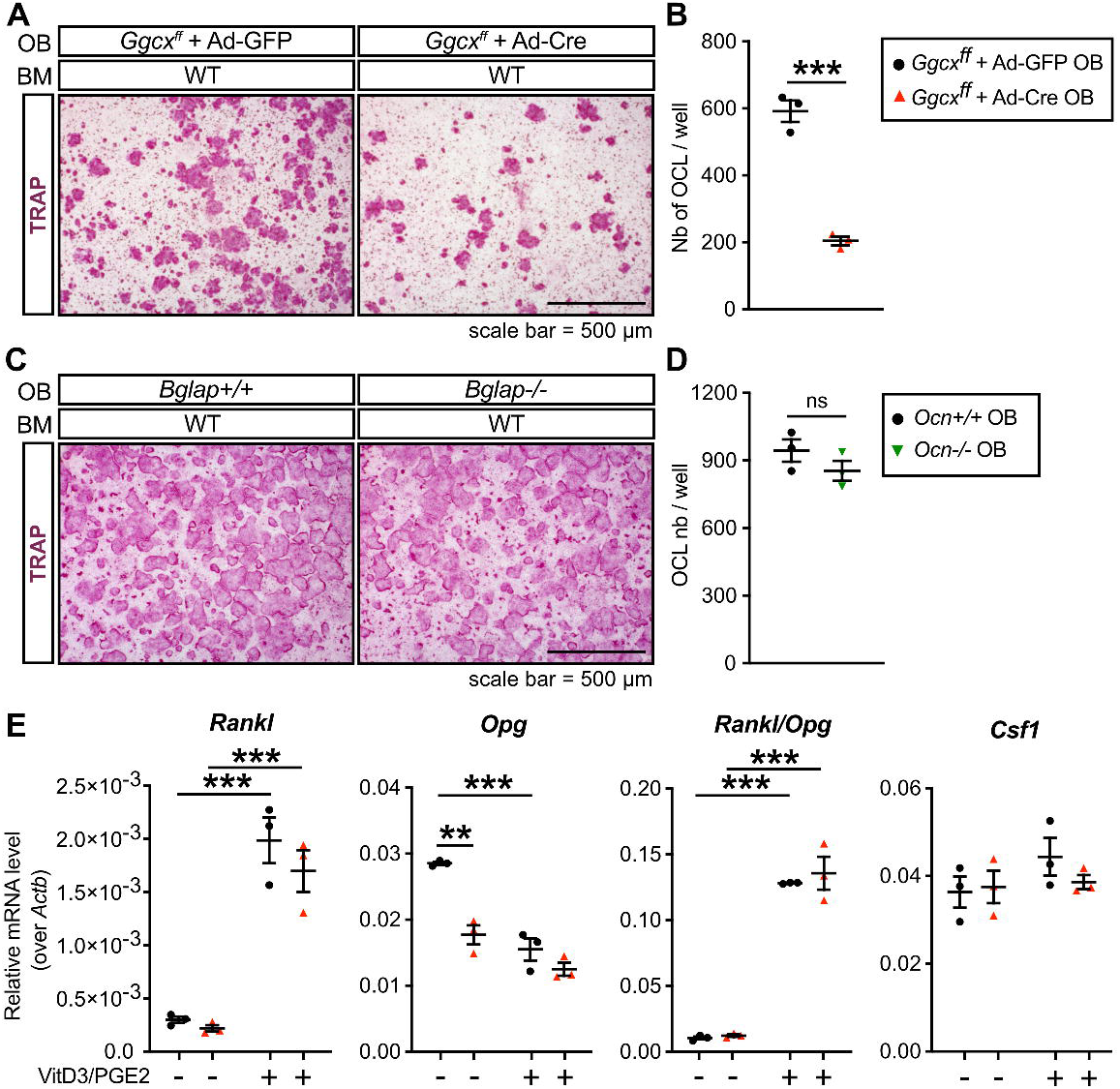
*Ggcx* inactivation impairs the ability of osteoblasts to support osteoclastogenesis ex vivo. **(A)** Representative TRAP staining of osteoblasts (OB) and bone marrow cells (BM) co-cultures at day 8 in the presence of prostaglandin E_2_ (PGE_2_; 10^-6^ M) and 1,25 vitamin D_3_ (VitD_3_; 10^-8^ M). *Ggcx^ff^* osteoblasts were transduced with either Ad-GFP (control) or Ad-Cre (knockout) before the addition of the WT bone marrow cells. **(B)** Quantification of the number of TRAP+ osteoclasts per well (Nb of OCL/well) (n=3). **(C)** Representative TRAP staining of osteoblasts and bone marrow cells co-cultures at day 8. Control (*Ocn+/+*) or osteocalcin-deficient (*Ocn-/-*) osteoblasts were cultured with WT bone marrow cells. **(D)** Quantification of the number of TRAP+ osteoclasts per well (Nb of OCL/well) (n=3). **(E)** Gene expression analysis by qPCR in *Ggcx^ff^*+ Ad-GFP and *Ggcx^ff^* + Ad-Cre osteoblasts cultured in presence (+) or absence (-) of PGE_2_ and VitD_3_ for 6 days. Results represent the mean ± SEM. Unpaired, 2-tailed Student’s t test was used in (B) and (D). Two-way ANOVA with Bonferroni’s posttests was used in (E). ***p < 0.001, **p < 0.01, *p < 0.05, ns: non-significant.

Cells from the osteoblast lineage produce factors that promote (i.e., RANKL and M-CSF) or inhibit (i.e., osteoprotegerin; OPG) osteoclast generation. We assessed the expression of these factors in control and GGCX-deficient osteoblasts to determine if γ-carboxylation regulates the pro-osteoclastogenic potential of osteoblasts. qPCR revealed that as expected, VitD3 and PGE2, robustly increased the expression of *Rankl* and suppressed *Opg* expression (**Fig. 3E**).

Consequently, the *Rankl/Opg* ratio, which is positively associated with osteoclast formation, was increased by more than tenfold in presence of VitD3 and PGE2. Importantly, the absence of *Ggcx* in osteoblasts had no significant impact on *Rankl, Opg* and the *Rankl/Opg* ratio in the pro-osteoclastogenic condition (**Fig. 3E**). In absence of VitD3 and PGE2, *Opg* expression was reduced (-38%) in GGCX-deficient osteoblasts, but this did not translate in a significant impact on the *Rankl/Opg* ratio, since the expression of *Rankl* was also decreased (-27%) in the same cells. The expression of *Csf1,* the gene encoding for M-CSF, was also unaffected in the absence of GGCX in osteoblasts (**Fig. 3E**). These data suggest that γ-carboxylation positively regulates osteoclast formation independently of osteocalcin and of the expression of known pro-osteoclastogenic factors.

### The γ-carboxylated protein GAS6 is expressed and secreted by osteoblasts and activates its receptors on pre-osteoclasts

Based on the results obtained in vivo and in co-culture assays, we postulated the existence of a γ-carboxylated protein, different from osteocalcin, secreted by osteoblasts and promoting osteoclast formation. Expression of all known γ-carboxylated protein encoding genes was measured by qPCR in primary mouse osteoblasts. As expected, we detected expression of the γ-carboxylated ECM proteins osteocalcin (*Ocn*), matrix Gla protein (*Mgp*) and periostin (*Postn*), previously shown to be produced by the osteoblastic lineage (**Fig.4A**) ^34–36^. Earlier studies have established that inactivation of *Mgp* or *Postn* in mice had no impact on osteoclast number and surface ^37,38^, and our own data show that osteocalcin is not required for osteoclastogenesis in coculture assays (**Fig.3C-D**). We therefore excluded these three proteins as potential mediators of the positive effect of γ-carboxylation on osteoclast formation. Of note, the γ-carboxylated coagulation factors and the Proline Rich and Gla domain 1 to 4 proteins (*Prrg1-4*) were undetectable in osteoblasts. However, the gene encoding Growth Arrest Specific 6 (*Gas6*) was expressed in osteoblasts and detected in the supernatant of osteoblasts cultured in pro-osteoclastogenic conditions (**Fig. 4A** and **Supplementary Fig. 2A**).

**Figure 4.**
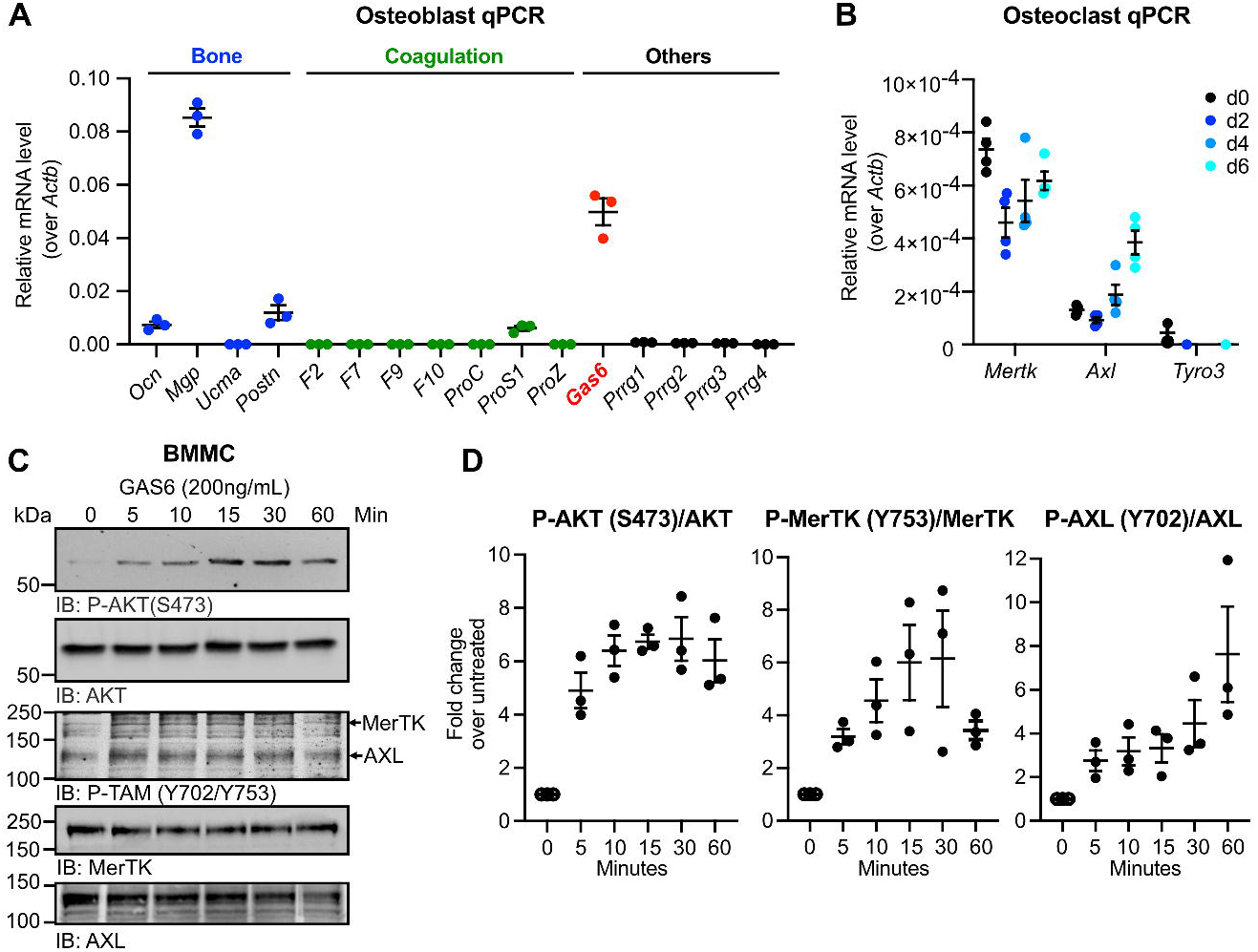
The γ-carboxylated protein GAS6 is expressed by osteoblasts and activates its receptors on pre-osteoclasts. **(A)** Expression analysis by qPCR of genes encoding the known γ-carboxylated proteins. *Gas6* is highlighted in red. **(B)** Gene expression analysis by qPCR of the TAM receptors *Axl*, *Mertk* and *Tyro3* in bone marrow derived monocytes (d0: day 0) and in differentiating osteoclast cultures in the presence of RANKL and M-CSF (d2-d6: day 2 to 6; n=4). **(C)** Western blot analysis of the phosphorylation (P) of AKT (S473) and TAM (Y702 in AXL and Y753 in MerTK) in bone marrow derived monocytes (BMMC) serum starved for 3h and treated with GAS6 (200ng/mL) for the indicated times. Total AKT, MerTK and AXL were used as loading controls. **(D)** Quantification of P-AKT (S473), P-MerTK (Y753) and P-AXL (Y702) normalized to the amount of total protein (n=3). Results represent the mean ± SEM.

GAS6 is a secreted γ-carboxylated protein that functions as a ligand for the TAM family of tyrosine kinase receptors that includes AXL, Tyro3 and MerTK. Given that GAS6 is a signaling molecule and that TAM receptors were previously shown to be present on myeloid cells in other tissues ^39^, we assessed the presence of functional GAS6 receptors at different stages of bone marrow-derived monocytes (BMMC) osteoclast differentiation. Both *MerTK* and *Axl* were detected in primary BMMC (d0) and in differentiating osteoclasts (d2, d4 and d6), while *Tyro3* expression was below the detection limit in most samples (**Fig.4B**).

We previously established that purified recombinant GAS6 (recGAS6) produced by HEK293 cells in presence of VK is fully γ-carboxylated and able to activate TAM receptors on muscle cells (**Supplementary Fig. 2B**) ^40^. We thus tested if this ligand could also elicit TAM receptors activation in primary pre-osteoclasts. For this purpose, BMMC were serum starved for 3 hours, stimulated for various times with recGAS6 (200ng/mL) and the phosphorylation of AXL and MerTK or their downstream target AKT ^41^ assessed by Western blotting. Following recGAS6 stimulation, an increase in the phosphorylation of MerTK (Y753) and AXL (Y702) was detected (**Fig.4C-D**). This tyrosine residue was previously shown to be a dominant autophosphorylation site in the tyrosine kinase domain of TAM receptors, which leads to the activation of their kinase activity ^42^. RecGAS6 also robustly induces the phosphorylation of AKT on serine 473, reflecting the activation of the phosphatidylinositol 3-kinase (PI3K) pathway (**Fig.4C-D**). Together, these results support the existence of a functional GAS6-TAM signaling axis coupling osteoblasts to osteoclasts.

### GAS6 signaling promotes osteoclast formation ex vivo

We then tested if GAS6-TAM signaling influences osteoclast formation using two approaches. In a first set of experiments, we assessed osteoclastogenesis in co-cultures of WT bone marrow cells and WT osteoblasts in the presence of increasing concentrations of two pharmacological TAM inhibitors, LDC1267 and R428 (bemcentinib) ^43,44^. As shown in Figure 5A, LDC1267, a pan-TAM inhibitor, dose dependently inhibits the formation of TRAP-positive multinucleated osteoclasts in these co-cultures, with an IC50 of ∼750nM. R428, which is >50-100-fold more selective for AXL than MerTK and Tyro3, also inhibits osteoclast formation with a lower IC50 of ∼250nM (**Fig.5B**). These data suggest that TAM receptor signaling promotes osteoclast formation in co-culture assay.

**Figure 5.**
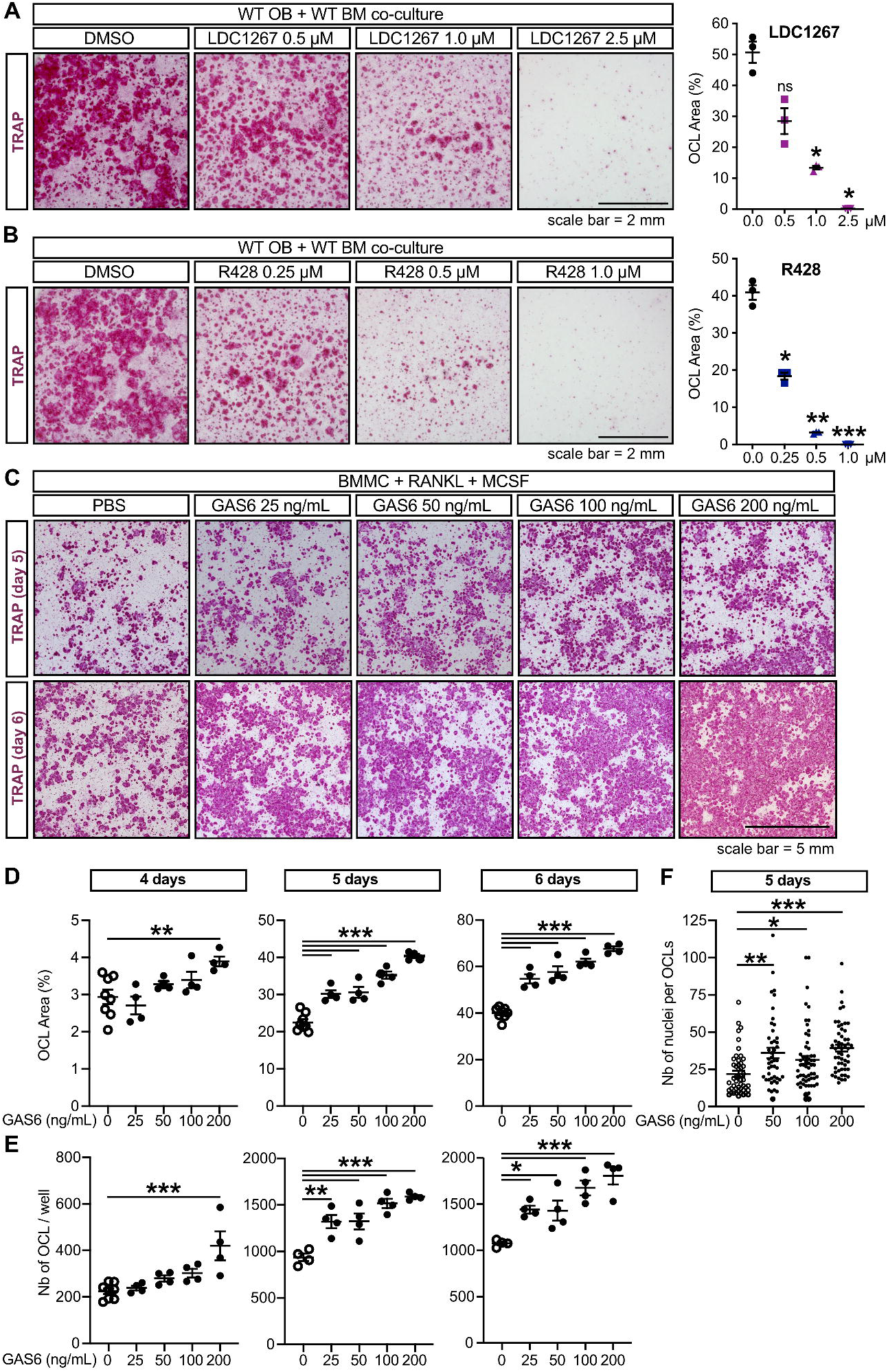
Carboxylated GAS6 signaling promotes osteoclast formation in culture. (A-B) Representative TRAP staining (left) and quantification of the TRAP+ osteoclast area (right) in WT osteoblast (OB) and bone marrow cell (BM) co-cultures at day 8 in the presence of PGE_2_ and VitD_3_, with or without the TAM inhibitors LDC1267 (A) or R428 (B), at the indicated concentrations. **(C-F)** Bone marrow derived monocytes (BMMC) were cultured in the presence of RANKL (20 ng/mL) and M-CSF (10 ng/mL) with or without recombinant carboxylated GAS6 at the indicated concentrations for up to 6 days. **(C)** Representative TRAP staining at day 5 and 6 of differentiation. **(D)** Quantification of the TRAP+ osteoclast area at day 4, 5 and 6 of differentiation. **(E)** Quantification of the number of TRAP+ multinucleated osteoclasts at day 4, 5 and 6 of differentiation. **(F)** Quantification of the number of nuclei per osteoclast at day 5 of differentiation. Results represent the mean ± SEM. One-way ANOVA with Bonferroni’s posttests was used in (A-B) and (D-F). ***p < 0.001, **p < 0.01, *p < 0.05, ns: non-significant.

In a second approach, we tested if exogenous recGAS6 (25-200 ng/mL) impacts osteoclast formation in primary BMMC cultures in the presence of RANKL (20ng/mL) and M-CSF (10ng/mL). In these experiments, recGAS6 significantly and dose-dependently increases osteoclast formation (**Fig.5C-E**). Quantification of the cultures following TRAP staining indicated that recGAS6 increased both the surface covered by TRAP+ osteoclasts (**Fig.5D**) and the number of multinucleated osteoclasts (**Fig. 5E)**, as early as 4 days of culture when small multinucleated TRAP-positive osteoclasts begin to appear. In addition, the number of individual nuclei per osteoclast was significantly increased in presence of recGAS6 (**Fig.5F**), indicating the formation of larger osteoclasts. Finally, supplementation of co-cultures of *Ggcx*-deficient osteoblasts and wild-type bone marrow cells with recGAS6 restored osteoclast numbers to levels comparable to controls (**Supplementary Fig. 3A-C**). These findings demonstrate that GAS6 is sufficient to rescue the impaired osteoclastogenesis caused by *Ggcx* deletion and identify GAS6-TAM signaling as a modulator of osteoclastogenesis ex vivo.

### GAS6 increases the fusion of pre-osteoclasts in culture

Osteoclast formation involves first, the transcriptional activation of a differentiation program in mononucleated pre-osteoclasts and second, their fusion to form mature multinucleated osteoclasts. We sought to determine if GAS6 signaling increases osteoclast formation by promoting differentiation of BMMC into osteoclasts. The expression of genes encoding several markers of osteoclast differentiation was measured by qPCR at 2, 4 and 6 days of differentiation in the absence or presence of intermediate doses of recGAS6 (i.e., 50 and 100 ng/mL). The presence of recGAS6 in the media did not significantly impact the expression of *Acp5* (TRAP) and *Clcn7* which gradually increase between 2 and 6 days of differentiation as expected (**Fig.6A-B**). The highest dose of GAS6 (100ng/mL) induces a small and significant increase in the expression of *Ctsk*, the gene encoding cathepsin K, at day 4 and 6 of differentiation, and of *Dcstamp* at day 6 of differentiation (**Fig.6C-D**). Overall, this marginal impact of GAS6 on the expression of the osteoclast differentiation program cannot explain its strong positive effect on the generation of multinucleated osteoclasts in the same culture conditions. Indeed, 50 and 100 ng/mL of GAS6 increases by >50% the number of osteoclasts and the area of TRAP+ osteoclasts (**Fig.5C-E**).

**Figure 6.**
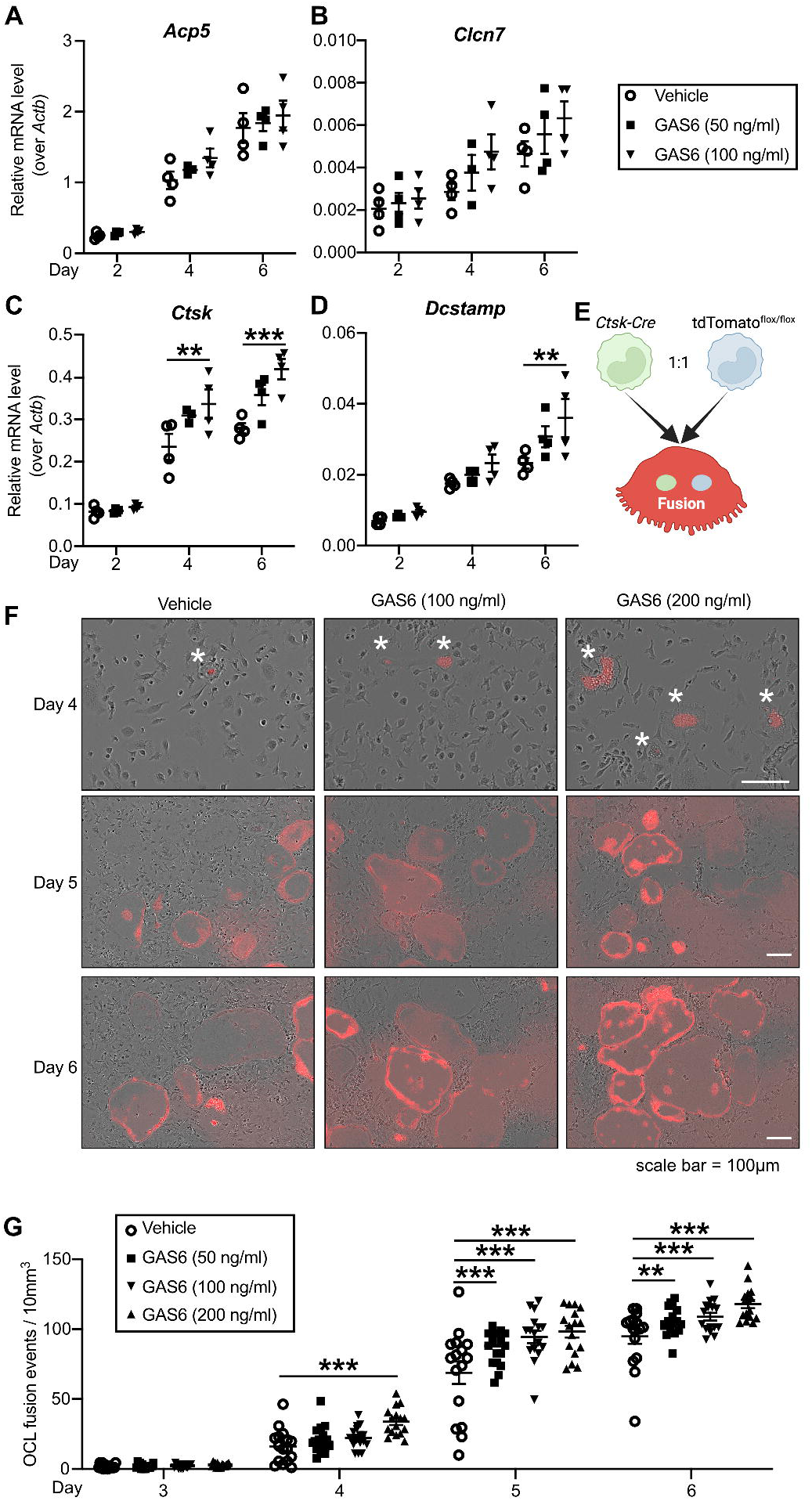
Carboxylated GAS6 impacts on osteoclast differentiation and fusion. (A-D) Gene expression analysis by qPCR of osteoclast differentiation markers *Acp5* (TRAP), *Clcn7*, *Ctsk* and *Dcstamp*. Bone marrow derived monocytes (BMMC) were cultured in the presence of RANKL (20 ng/mL) and M-CSF (10 ng/mL) with or without recombinant carboxylated GAS6 at the indicated concentrations for 2, 4 and 6 days (n=4 per condition). **(E)** Schematic representation of the assay used to assess the impact of GAS6 on pre-osteoclast fusion in culture using a conditionally activated tdTomato (Tom) reporter. **(F)** Representative pictures of live osteoclast cultures at the indicated time and concentration of recombinant GAS6. The stars indicate the presence of fusion events (Tom+ cells) in the GAS6 conditions at Day 4. **(G)** Quantification of the number of fusion events per 10 mm^2^ at the indicated time of osteoclasts differentiation (n=16 fields per condition). Results represent the mean ± SEM. One-way ANOVA with Bonferroni’s posttests was used in (A-D) and (G). ***p < 0.001, **p < 0.01.

Since recGAS6 increases the number of nuclei per OCL (**Fig.5F**), we next tested if GAS6 promotes pre-osteoclast fusion. For this purpose, we established a pre-osteoclast fusion assay in which an equal number of *Ctsk^Cre/+^*and *Rosa26^CAG-lox-stop-lox-tdTomato^*BMMCs were induced to differentiate in osteoclasts. In this experimental setting, the fluorescent protein tdTomato (Tom) is only expressed in cells resulting from the fusion of at least one *Ctsk^Cre/+^* and one *Rosa26^CAG-lox-stop-lox-tdTomato^* pre-osteoclast (**Fig.6E**). We then monitored using time-lapse microscopy the appearance of fluorescent Tom+ cells in the presence or absence of recGAS6. As shown in Figure 6F, Tom+ cells begin to appear in osteoclast cultures at day 4 and increased in number considerably at day 5 and 6, validating our approach. Interestingly, the number of Tom+ cells was significantly higher in the presence of recGAS6 in the culture media even with the lowest dose tested (50 ng/mL), suggesting that GAS6 signaling promotes osteoclast fusion.

### GAS6 promotes osteoclast formation and bone resorption in vivo

In addition to GAS6, protein S (Pros1) can also activate TAM receptors ^45^. Our data show that osteoblasts express Pros1 (**Fig. 4A**) and pre-osteoclasts express both AXL and MerTK (**Fig. 4B**). This redundancy in the TAM signaling system prevented us from assessing its role in osteoclast formation using a simple genetic loss-of-function model. Therefore, we employed a gain-of-function approach to evaluate the effect of increased GAS6 on osteoclastic bone resorption in vivo, using a previously described transgenic mouse line that expresses GAS6 under the human *ApoE* promoter ^40^. In these *ApoE-Gas6^Tg^* mice, GAS6 is produced by hepatocytes, fully carboxylated and secreted in the circulation, resulting in a fourfold increase in active GAS6 in serum (**Fig.7A**). Because the skeleton, particularly the bone marrow cavity, is highly vascularized, we reasoned that this rise in circulating GAS6 should lead to higher concentration in the bone marrow, where osteoclast differentiation occurs. Consistent with this, GAS6 levels in the bone marrow were doubled in *ApoE-Gas6^Tg^* mice (**Fig.7B**). We then examined bone density and remodeling in this GAS6 gain-of-function model.

**Figure 7.**
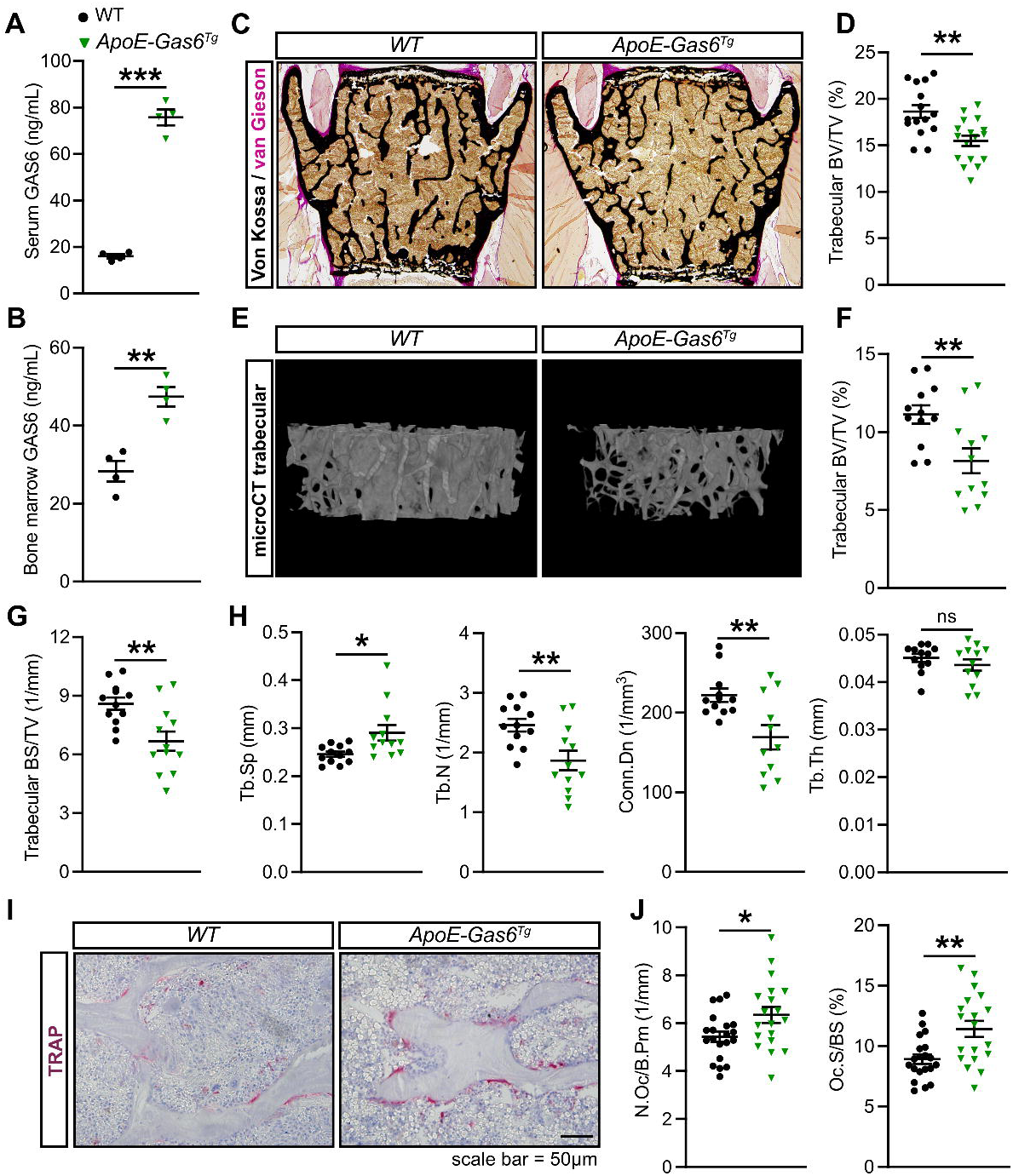
GAS6 promotes osteoclast formation and bone resorption in vivo. (A-J) Six-month-old *WT* (non-transgenic littermates) and *ApoE-Gas6^Tg^* male mice were analyzed. **(A-B)** GAS6 concentration in the serum (A) and bone marrow cavity (B) (n=4). **(C)** Representative pictures of sections from lumbar vertebrae stained with von Kossa and van Gieson. **(D)** Quantification of trabecular bone volume over tissue volume (BV/TV) from the L4 and L5 lumbar vertebrae sections (n=15-17). **(E-H)** μCT analysis of the distal femur trabecular bone (n=12). **(E)** Representative μCT images. **(F)** Quantification of trabecular bone volume (BV/TV). **(G)** Quantification of trabecular bone surface density (BS/TV). **(H)** Trabecular bone µCT derived data. Tb.Sp., Tb.N., and Tb.Th., trabecular spacing, number, and thickness, respectively; Conn.Dn., connectivity density. **(I-J)** Bone histomorphometry analysis of lumbar vertebrae in six-month-old *WT* and *ApoE-Gas6^Tg^* male (n=19-20). **(I)** Representative pictures of TRAP staining. **(J)** Number of osteoclasts per bone perimeter (N.Oc/B.Pm) and osteoclast surface over bone surface (Oc.S/BS). Results represent the mean ± SEM. Unpaired, 2-tailed Student’s t test was used in (A, B, D, F, G and J). ***p < 0.001, **p < 0.01, *p < 0.05, ns: non-significant.

Histology and µCT revealed a reduced trabecular bone volume (BV/TV) in vertebrae and femur of 6-month-old *ApoE-Gas6^Tg^* male mice compared to WT littermates (**Fig.7C-F**). The µCT analyses showed that trabecular bone surface density (BS/TV) was also significantly reduced in the same animals (**Fig. 7G**). Trabecular spacing (Tb.Sp) was significantly increased and trabecular number (Tb.N) and connectivity density (Conn.Dn) significantly decreased, while trabecular thickness (Tb.Th) was unaffected in *ApoE-Gas6^Tg^* male mice (**Fig. 7H**). Static and dynamic bone histomorphometry indicated that *ApoE-Gas6^Tg^* male mice were characterized by a significant increase in both the number (N.Oc/B.Pm) and surface (Oc.S/BS) of osteoclasts in the trabecular bone (**Fig. 7I-J**). Importantly, bone formation parameters were unaffected in the same animals (**Supplementary Fig. 4**). Altogether, these data indicate that an increased concentration of GAS6 in the bone marrow is sufficient to stimulate the formation of osteoclasts in vivo, resulting in reduced bone density.

## Discussion

Whether VK supplementation has any beneficial impact on skeletal health remains unclear since interventional studies in humans have led to conflicting results ^22^. Our data suggest that VK-dependent carboxylation in osteoblasts is not required for normal bone formation and mineralization by this cell type. Instead, we uncover that γ-carboxylation, through GAS6, impacts the capacity of osteoblasts to regulate osteoclast formation. Taken at face value, these findings suggest that VK should negatively regulate bone mass, potentially explaining the limited impact of VK supplementation on bone density in humans. Conversely, osteoclastic bone resorption also plays a critical role in bone turnover and repair of microdamages to the bone structure ^46^.

Although the potential role of γ-carboxylation or GAS6 in bone regeneration was not formally tested in the current study, it is possible that the production of active γ-carboxylated GAS6 by osteoblasts may be important for the fine tuning of osteoclastic activity during bone repair processes, thereby promoting the maintenance of a healthy skeleton and preventing fractures. Consistent with this, a meta-analysis of several large observational studies found that in individuals with atrial fibrillation, treatment with warfarin, an inhibitor of VKORC1, significantly increases the risk of hip and vertebral fractures compared with direct oral anticoagulants that do not target carboxylation ^47^.

Vitamin K-dependent carboxylation in osteoblasts may influence the activity of several Gla proteins, including osteocalcin and MGP. The conclusion that γ-carboxylation in osteoblasts impact bone resorption through GAS6 is supported by several lines of evidence. First, the bone phenotype of the *ApoE-Gas6^Tg^* mice, a GAS6 gain-of-function model, is the mirror image of the one observed in *Ggcx^fl/fl^*;*OCN-Cre* mice. Second, in line with previous reports, we show that GAS6 is strongly expressed in osteoblasts and secreted by this cell type ^48,49^. Third, recombinant carboxylated GAS6 promotes the formation of larger osteoclasts in culture, while pharmacological inhibition of TAM receptors reduces osteoclastogenesis. Fourth, osteocalcin-deficient osteoblasts fully support osteoclast formation, while *Mgp*-deficient mice have normal bone resorption ^38^. Of note, mineral apposition rate (MAR) was reduced in *Ggcx^fl/fl^*;*OCN-Cre* mice, but unchanged in *ApoE-Gas6^Tg^* mice, suggesting that additional, yet to be identified Gla protein(s), may be involved in regulating bone formation.

In line with our findings, a recent study reported that whole body inactivation of MerTK in mice results in higher bone density with concomitant reduction in bone resorption ^50^. TAM receptors signaling may also play a role in osteoblasts function since it was reported that MerTK inactivation specifically in osteoblasts using the *Col1a1*-*Cre* driver caused increased bone formation and bone density, without affecting bone resorption ^49^. Importantly, *Ggcx^fl/fl^*;*OCN-Cre* mice did not show an increase in bone formation, suggesting that osteoblasts-derived carboxylated GAS6 is not involved in suppressing osteoblast activity through MerTK. Both protein S and GAS6 can activate MerTK, while AXL is activated exclusively by GAS6 ^45^.

According to our own data and previous reports ^49,51^, osteoblasts express protein S. We also found that pre-osteoclasts and differentiated osteoclasts express comparable level of AXL and MerTK. Due to this redundancy in ligands and receptors, assessing the role of the Pros1/GAS6/TAM axis in bone resorption using loss-of-function mouse models will likely require the generation of cell-specific double knockout mice.

In our primary culture assays, low to intermediate doses of GAS6 (i.e., 25 to 100 ng/mL) significantly increased osteoclasts number and surface at 5 and 6 days of culture. The same concentration also increased the average number of nuclei per osteoclast. However, dose of 50 to 100 ng/mL had limited impact on the expression of the osteoclast differentiation program.

Together with the results of our fusion assay, these data suggest that GAS6 does not regulate osteoclast differentiation but rather promotes the fusion of pre-osteoclasts.

To fully activate TAM receptors, GAS6 requires its Gla domain, even though this domain is not necessary for binding to the TAM ectodomain ^45^. The GAS6 Gla domain can bind to phosphatidylserines (PtdSer) exposed on the surface of apoptotic cells and stimulate phagocytosis by TAM receptor-expressing myeloid cells. Although the precise mechanism remains unclear, PtdSer binding induces a conformational change in GAS6 that enables TAM receptor activation on phagocytic cells ^52^. Recent studies have shown that PtdSer are exposed on the outer leaflet of the plasma membrane during pre-osteoclast differentiation, and this exposure facilitates the fusion of osteoclast precursors ^53–55^. Therefore, GAS6 may promote this fusion process by binding PtdSer on one pre-osteoclast while simultaneously activating TAM receptors on another one. Future work will aim at elucidating how TAM signaling in pre-osteoclasts contributes to cell fusion.

This study has some limitations. First, in vivo analyses by μCT and histology were performed exclusively in male mice. Although this was partially addressed by using female bone marrow cells in ex vivo experiments, sex-specific effects cannot be ruled out. Second, the impact of GAS6 and γ-carboxylation on human osteoclast formation was not evaluated. Finally, we cannot exclude the possibility that GAS6 is not the only γ-carboxylated protein mediating the effects of vitamin K on bone cells.

In conclusion, this work uncovers a new vitamin K-dependent carboxylation pathway in osteoblasts involved in the regulation of osteoclast formation.

## Materials and Methods

### Animals

All the mouse strains used in this study were backcrossed or generated on a pure C57BL6/J genetic background. The generation of *Ggcx^fl/fl^*;*OCN-Cre* mice was previously described ^33^. Briefly, *Ggcx^flox/flox^* mice were bred with *Osteocalcin-Cre* (OCN-Cre) mice, which express the Cre recombinase in mature osteoblasts only ^32^. We previously established that *Ggcx* was specifically and efficiently inactivated in osteoblasts in *Ggcx^fl/fl^*;*OCN-Cre* mice ^33^. The generation and characterization of *ApoE-Gas6^Tg^* mice expressing mouse *Gas6* under the control of the liver specific human APOE promoter and hepatic enhancers was also previously reported ^40^. The *tdTomato^flox/flox^* strain expressing the tdTomato reporter gene from the *Rosa26* locus was obtained from the Jackson Laboratory (*B6.Cg-Gt(ROSA)26Sor^tm^*^14^*^(CAG-tdTomato)Hze/J^*; stock 007914). The *Ctsk-Cre* (*Ctsk^tm1(cre)Ska^*) strain was previously described ^56^. Osteocalcin-deficient mice (*Ocn-/-*) in which both osteocalcin genes (*Bglap1/Ocn1* and *Bglap2/Ocn2*) are inactivated were previously reported ^29^. Male mice were used in all in vivo experiments. C57BL6/J male and female newborns or female adult mice (Jackson Laboratory; strain 000664) were used as a source of primary wildtype (WT) osteoblasts and bone marrow cells respectively. All strains were maintained in an IRCM specific pathogen-free animal facility under 12-hour dark/12-hour light cycles. Mice were fed ad libitum a normal chow diet (Teklad global 18% protein rodent diet; 2918; Envigo; containing 50mg/kg of VK_3_ and ∼100μg/kg of VK_1_). All animal use complied with the guideline of the Canadian Committee for Animal Protection (CCAP) and was approved by IRCM institutional animal care committee.

### Biochemical measurements

Serum carboxylated (Gla) and total osteocalcin were assessed using previously described ELISA assays ^57^. Serum C-terminal telopeptide of collagen type I (CTX) was measured using a commercially available assay (RatLaps; IDS, AC-06F1). GAS6 from serum and osteoblast supernatant were quantified using a mouse GAS6 ELISA (DuoSet ELISA, R&D Systems, DY986) as we previously described ^40^. To measure GAS6 in bone marrow, the bone marrow of one femur per mouse was flushed using 1mL of PBS 1× and the concentration of GAS6 quantified using the same ELISA assay. To estimate the concentration of GAS6 in the BM, the amount of GAS6 per femur (i.e., GAS6 concentration × 1mL) was divided by 20μL, the estimated volume for a femur BM cavity of a 6-month-old male mouse based on previous μCT data.

### Bone histology and histomorphometry

Mice were injected intraperitoneally twice at a 3-day interval with 0.2 mL of a solution of calcein (0.25% in a solution of 0.15M NaCl and 2% NaHCO_3_). Following euthanasia, the skeletal tissue was fixed in phosphate buffered 4% formaldehyde for 24h, followed by 24h dehydration in 70% ethanol, and processed for histomorphometry and micro-CT. For non-decalcified histology, the vertebrae were embedded in methyl methacrylate resin (MMA) as previously described ^58^ and sectioned (5 μm and 7μm thickness). Following deplastification, the sections were stained with von Kossa and van Gieson (7μm), Toluidine blue (5 μm) or TRAP (5 μm). Bone histomorphometric analyses were completed using the OsteoMeasure Analysis System (OsteoMetrics) connected to a light and fluorescent microscope (DM4000B LED; Leica) with a 40× objective (HCX PL FLUOTAR 40×; NA = 0.75).

### Micro-CT

Ex vivo micro-computed tomography (μCT) scans were performed on harvested distal femurs using the Skyscan 1272 system (Bruker, Kontich, Belgium) at 62 kV, 156 µA, and 6 µm isotropic resolution with a 0.5 mm aluminum filter. Images were reconstructed with NRecon (beam hardening correction: 50%; ring artifact reduction: 7) and analyzed using CTAn software (Bruker, Skyscan). The trabecular bone region of interest (ROI) was defined 0.75 mm proximal to the growth plate and extended 1 mm proximally. Morphometric parameters were calculated following published guidelines for rodent bone analysis using μCT ^59^.

### Production and Purification of Recombinant GAS6

Recombinant γ-carboxylated mouse GAS6 was produced as previously described ^40^. Briefly, HEK293 Flp-In T-REx cells (Thermo Fisher Scientific) stably transfected with a tetracycline-inducible plasmid encoding mouse GAS6 with a C-terminal 6xHis tag were maintained in DMEM (Wisent) supplemented with 10% fetal bovine serum (FBS; Sigma) and 200 μg/mL hygromycin for selection. Cells were expanded and the expression of GAS6 induced with 1 μg/mL tetracycline in presence of 22 μM vitamin K_1_ (Millipore Sigma, V3501) in serum-free DMEM for 36 hours. The culture supernatant was collected, centrifuged, and filtered (0.22 μm) to remove cells and debris. Recombinant GAS6 was purified by nickel affinity chromatography using a HisTrap HP column (Cytiva) connected to an ÄKTAprime FPLC system (GE Healthcare). Elution was performed using an imidazole-containing buffer, and the purified protein was dialyzed against PBS 1‘. Protein purity and concentration were assessed by SDS-PAGE followed by Coomassie staining (see **Supplementary Fig. 2B**).

### Osteoblast cultures

Mouse calvaria pre-osteoblasts were isolated from 3-day-old male and female pups and cultured as previously described ^33^. Briefly, dissected calvarias were digested at 37°C twice for 10 minutes and twice for 30 min with a solution of collagenase type 2 (0.1 mg/mL; Worthington Biochemical Corporation, LS004176) containing 0.25% trypsin. Only the last two digestions were kept and cultured in alpha modified minimum essential medium (α-MEM, Sigma, M0644) supplemented with 10% FBS and 1% penicillin-streptomycin (Wisent) to obtain proliferating pre-osteoblasts (pre-OB). Differentiation into mineralizing osteoblasts (OB) was induced by culturing these cells in the same media supplemented with 5mM b-glycerophosphate and 100 mg/mL L-ascorbic acid for 21 days. Media was changed every 2 days. To generate control and *Ggcx^-/-^* pre-osteoblasts ex vivo, *Ggcx^fl/fl^* cells were transduced with either GFP- or Cre-expressing adenovirus (University of Iowa) at a MOI of 200 ^33^.

### Bone marrow monocytes and osteoclast cultures

Primary bone marrow cells were isolated from the femurs and tibias of 6- to 10-week-old C57BL6/J female mice. Bones were cleaned of soft tissue and flushed with α-MEM supplemented with 10% FBS and 1% penicillin-streptomycin using a 25G needle. Bone marrow-derived monocytes (BMMC) were obtained by culturing the marrow cell suspension in non-adherent 10 cm dishes in M-CSF–rich medium (20% L929 cell-conditioned medium; ATCC) for 5 days. Non-adherent cells were removed by vigorous washing with PBS 1‘, and adherent cells were harvested by incubation with 0.02% EDTA in PBS 1‘. In some experiments, BMMC were starved for 3 h in serum-free medium supplemented with 10 mM HEPES (pH 7.4) and 0.1% BSA and stimulated with 200 ng/mL of recombinant carboxylated GAS6 for 5 to 60 minutes.

For osteoclast-like cell (OCL) differentiation, BMMCs were seeded at 2 × 10^4^ cells per well in 24-well plates and cultured in medium supplemented with 20 ng/mL RANKL and 10 ng/mL M-CSF, with or without recombinant γ-carboxylated GAS6 at various concentrations (25-200 ng/mL). Multinucleated mature osteoclasts were visualized by TRAP staining ^60^. In selected experiments, nuclei were stained with DAPI, and both brightfield and fluorescence images were acquired (DM4000B LED; Leica) with a 5× objective (HCX PL FLUOTAR 5×; AP = 0.15). The number of nuclei per osteoclast was quantified using ImageJ. In co-culture assays, 4.5 × 10^6^ bone marrow cells were added to confluent osteoblast cultures in 12 well plates in αMEM supplemented with 10% FBS, prostaglandin E_2_ (PGE_2_; 10^-6^ M) and 1,25 vitamin D_3_ (VitD_3_; 10^-8^ M). In some experiments, TAM receptor inhibitors R428 or LDC1267 (Selleckchem, S2841 and S7638) were added to the cultures at the indicated concentrations. After 7 to 8 days of co-culture, the cells were fixed and TRAP staining performed. Osteoclast culture pictures were acquired using a SteREO Discovery.V12 microscope (Zeiss) using Zen software. Osteoclast number and surface area were quantified from TRAP-stained cell cultures using ImageJ software.

### Fusion assay

An equal number of *tdTomato^flox/flox^*and *Ctsk-Cre* BMMCs (1 × 10^4^ cells each per well) were seeded in 24-well plates and cultured in medium supplemented with 20 ng/mL RANKL and 10 ng/mL M-CSF, with or without recombinant γ-carboxylated GAS6 at various concentrations.

Plates were imaged using an Incucyte Live-Cell Analysis System (Sartorius). Fluorescence (tdTomato) and phase contrast images were acquired every 24 h starting from day 3 of culture using a 10× objective (16 fields per well). Images were analyzed using ImageJ and each tdTomato-positive (red) cell with 2 or more nuclei was counted as a fusion event.

### Gene expression

Total RNA was extracted from cells or tissues as previously ^61^. mRNA was reverse transcribed using M-MLV as previously reported ^62^. Relative gene expression was measured by quantitative PCR using PowerUp SYBR Green Master Mix (A25741; Applied Biosystems) with gene-specific primers (Supplemental Table 1) on a ViiA7 Real-Time PCR System (Applied Biosystems).

### Western blot

Proteins from cells or tissues were extracted with a lysis buffer containing 20mM Tris-HCl (pH 7.5), 150mM NaCl, 1mM EDTA (pH 8.0), 1mM EGTA, 2.5mM NaPyrophosphate, 1mM β-glycerophosphate, 10mM NaF, 1% Triton, 1 mM vanadate, 1mM phenylmethylsulfonyl fluoride (PMSF) and protease inhibitors (Roche Diagnostics, 4693132001). Proteins were detected by Western blot with the indicated primary antibodies overnight at 4°C and quantified by densitometry analyses using Image Lab software (Bio-Rad Laboratories). The antibodies used are listed in Supplemental Table 2.

## Statistical Analysis

Statistical analyses were performed with GraphPad Prism version 10.4.0. Results are given as mean ± SEM. Unpaired two-tailed Student’s *t* test was used to compare two groups. For experiments involving multiple groups, one-or two-way ANOVA, followed by Bonferroni’s multiple comparisons test was used. In all figures, ∗ *P* < 0.05; ∗∗ *P* < 0.01; ∗∗∗ *P* < 0.001. All experiments were repeated at least 3 times or performed on at least 3 independent animals.

## Data and Resource Availability

The data sets generated and/or analyzed during the current study are available from the corresponding author upon reasonable request. The *ApoE-Gas6*^Tg^ transgenic and the *Ggcx^flox/flox^* mouse lines, the recombinant GAS6 expressing cell line, the phosphorylated AXL (Y702) antibody and the mouse VKORC1 antibody used during the current study are available from the corresponding author upon reasonable request.

## Acknowledgements

We thank S. Kato and T. Clemens for providing *Ctsk-Cre* and *OCN-*Cre mice respectively. This work was supported by funding from the Fonds de Recherche du Québec–Santé (to M.F.), Canada Research Chairs program (to M.F.) and Canadian Institutes of Health Research grant PJT-159534 (to M.F.). B.A.R. received a postdoctoral fellowship from the Fonds de Recherche du Québec–Santé.

## Contributions

M.F. and M.P. assembled the figures. M.F. wrote the manuscript with suggestions from the other authors. M.P., M.F., D.N.T.P., J.L., B.A.R. and Y.W.K., collected and analyzed data. M.F. conceived and designed the project and provided supervision to M.P., D.N.T.P., J.L., B.A.R. and Y.W.K. μCT data were collected and analyzed by A.G.A.A. and M.M. M.F. is the guarantor of this work and, as such, had full access to all the data in the study and takes responsibility for the integrity of data and the accuracy of data analysis.

## References

1 Morris, J. A. et al. An atlas of genetic influences on osteoporosis in humans and mice. Nat Genet 51, 258–266, doi:10.1038/s41588-018-0302-x (2019).

2 Zhu, X., Bai, W. & Zheng, H. Twelve years of GWAS discoveries for osteoporosis and related traits: advances, challenges and applications. Bone Res 9, 23, doi:10.1038/s41413-021-00143-3 (2021).

3 Moller, A. M. J. et al. Aging and menopause reprogram osteoclast precursors for aggressive bone resorption. Bone Res 8, 27, doi:10.1038/s41413-020-0102-7 (2020).

4 Wilson-Barnes, S. L., Lanham-New, S. A. & Lambert, H. Modifiable risk factors for bone health & fragility fractures. Best Pract Res Clin Rheumatol 36, 101758, doi:10.1016/j.berh.2022.101758 (2022).

5 Nieves, J. W. Osteoporosis: the role of micronutrients. Am J Clin Nutr 81, 1232S–1239S, doi:10.1093/ajcn/81.5.1232 (2005).

6 Veldurthy, V. et al. Vitamin D, calcium homeostasis and aging. Bone Res 4, 16041, doi:10.1038/boneres.2016.41 (2016).

7 Brzezinska, O., Lukasik, Z., Makowska, J. & Walczak, K. Role of Vitamin C in Osteoporosis Development and Treatment-A Literature Review. Nutrients 12, doi:10.3390/nu12082394 (2020).

8 Clements, M. et al. A 2-Year Randomized Controlled Trial With Low-Dose B-Vitamin Supplementation Shows Benefits on Bone Mineral Density in Adults With Lower B12 Status. J Bone Miner Res 37, 2443–2455, doi:10.1002/jbmr.4709 (2022).

9 Tucker, K. L. et al. Low plasma vitamin B12 is associated with lower BMD: the Framingham Osteoporosis Study. J Bone Miner Res 20, 152–158, doi:10.1359/JBMR.041018 (2005).

10 Booth, S. L. et al. Vitamin K intake and bone mineral density in women and men. Am J Clin Nutr 77, 512–516 (2003).

11 Booth, S. L. et al. Associations between vitamin K biochemical measures and bone mineral density in men and women. J Clin Endocrinol Metab 89, 4904–4909, doi:10.1210/jc.2003-031673 (2004).

12 Booth, S. L. et al. Dietary vitamin K intakes are associated with hip fracture but not with bone mineral density in elderly men and women. Am J Clin Nutr 71, 1201–1208, doi:10.1093/ajcn/71.5.1201 (2000).

13 Feskanich, D. et al. Vitamin K intake and hip fractures in women: a prospective study. Am J Clin Nutr 69, 74–79 (1999).

14 Hao, G. et al. Vitamin K intake and the risk of fractures: A meta-analysis. Medicine (Baltimore) 96, e6725, doi:10.1097/MD.0000000000006725 (2017).

15 Hamidi, M. S., Gajic-Veljanoski, O. & Cheung, A. M. Vitamin K and bone health. Journal of clinical densitometry: the official journal of the International Society for Clinical Densitometry 16, 409–413, doi:10.1016/j.jocd.2013.08.017 (2013).

16 Thijssen, H. H. & Drittij-Reijnders, M. J. Vitamin K distribution in rat tissues: dietary phylloquinone is a source of tissue menaquinone-4. The British journal of nutrition 72, 415–425 (1994).

17 Ronden, J. E., Thijssen, H. H. & Vermeer, C. Tissue distribution of K-vitamers under different nutritional regimens in the rat. Biochim Biophys Acta 1379, 16–22 (1998).

18 Fang, Y., Hu, C., Tao, X., Wan, Y. & Tao, F. Effect of vitamin K on bone mineral density: a meta-analysis of randomized controlled trials. J Bone Miner Metab 30, 60–68, doi:10.1007/s00774-011-0287-3 (2012).

19 Cockayne, S. et al. Vitamin K and the prevention of fractures: systematic review and meta-analysis of randomized controlled trials. Archives of internal medicine 166, 1256–1261, doi:10.1001/archinte.166.12.1256 (2006).

20 Inoue, T. et al. Randomized controlled study on the prevention of osteoporotic fractures (OF study): a phase IV clinical study of 15-mg menatetrenone capsules. J Bone Miner Metab 27, 66–75, doi:10.1007/s00774-008-0008-8 (2009).

21 Cheung, A. M. et al. Vitamin K supplementation in postmenopausal women with osteopenia (ECKO trial): a randomized controlled trial. PLoS medicine 5, e196, doi:10.1371/journal.pmed.0050196 (2008).

22 Dupuy, M. et al. Vitamin K: Metabolism, Genetic Influences, and Chronic Disease Outcomes. Food Sci Nutr 13, e70431, doi:10.1002/fsn3.70431 (2025).

23 Zhou, M., Han, S., Zhang, W. & Wu, D. Efficacy and safety of vitamin K2 for postmenopausal women with osteoporosis at a long-term follow-up: meta-analysis and systematic review. J Bone Miner Metab 40, 763–772, doi:10.1007/s00774-022-01342-6 (2022).

24 Salma et al. Effect of Vitamin K on Bone Mineral Density and Fracture Risk in Adults: Systematic Review and Meta-Analysis. Biomedicines 10, doi:10.3390/biomedicines10051048 (2022).

25 Lacombe, J. & Ferron, M. Vitamin K-dependent carboxylation in beta-cells and diabetes. Trends Endocrinol Metab 35, 661–673, doi:10.1016/j.tem.2024.02.006 (2024).

26 Berkner, K. L. & Runge, K. W. Vitamin K-Dependent Protein Activation: Normal Gamma-Glutamyl Carboxylation and Disruption in Disease. International journal of molecular sciences 23, doi:10.3390/ijms23105759 (2022).

27 Hauschka, P. V., Lian, J. B. & Gallop, P. M. Direct identification of the calcium-binding amino acid, gamma-carboxyglutamate, in mineralized tissue. Proc Natl Acad Sci U S A 72, 3925–3929 (1975).

28 Lambert, L. J. et al. Increased trabecular bone and improved biomechanics in an osteocalcin-null rat model created by CRISPR/Cas9 technology. Disease models & mechanisms 9, 1169–1179, doi:10.1242/dmm.025247 (2016).

29 Ducy, P. et al. Increased bone formation in osteocalcin-deficient mice. Nature 382, 448–452 (1996).

30 Diegel, C. R. et al. An osteocalcin-deficient mouse strain without endocrine abnormalities. PLoS genetics 16, e1008361, doi:10.1371/journal.pgen.1008361 (2020).

31 Moriishi, T. et al. Osteocalcin is necessary for the alignment of apatite crystallites, but not glucose metabolism, testosterone synthesis, or muscle mass. PLoS genetics 16, e1008586, doi:10.1371/journal.pgen.1008586 (2020).

32 Zhang, M. et al. Osteoblast-specific knockout of the insulin-like growth factor (IGF) receptor gene reveals an essential role of IGF signaling in bone matrix mineralization. J Biol Chem 277, 44005–44012 (2002).

33 Ferron, M., Lacombe, J., Germain, A., Oury, F. & Karsenty, G. GGCX and VKORC1 inhibit osteocalcin endocrine functions. J Cell Biol 208, 761–776, doi:10.1083/jcb.201409111 (2015).

34 Coutu, D. L. et al. Periostin, a member of a novel family of vitamin K-dependent proteins, is expressed by mesenchymal stromal cells. J Biol Chem 283, 17991–18001 (2008).

35 Barone, L. M. et al. Developmental expression and hormonal regulation of the rat matrix Gla protein (MGP) gene in chondrogenesis and osteogenesis. J Cell Biochem 46, 351–365, doi:10.1002/jcb.240460410 (1991).

36 Ducy, P. & Karsenty, G. Two distinct osteoblast-specific cis-acting elements control expression of a mouse osteocalcin gene. Mol Cell Biol 15, 1858–1869 (1995).

37 Gerbaix, M., Vico, L., Ferrari, S. L. & Bonnet, N. Periostin expression contributes to cortical bone loss during unloading. Bone 71, 94–100, doi:10.1016/j.bone.2014.10.011 (2015).

38 Marulanda, J., Gao, C., Roman, H., Henderson, J. E. & Murshed, M. Prevention of arterial calcification corrects the low bone mass phenotype in MGP-deficient mice. Bone 57, 499–508, doi:10.1016/j.bone.2013.08.021 (2013).

39 Lemke, G. Biology of the TAM receptors. Cold Spring Harb Perspect Biol 5, a009076, doi:10.1101/cshperspect.a009076 (2013).

40 Schott, C. et al. GAS6 and AXL Promote Insulin Resistance by Rewiring Insulin Signaling and Increasing Insulin Receptor Trafficking to Endosomes. Diabetes 73, 1648–1661, doi:10.2337/db23-0802 (2024).

41 Goruppi, S., Ruaro, E., Varnum, B. & Schneider, C. Requirement of phosphatidylinositol 3-kinase-dependent pathway and Src for Gas6-Axl mitogenic and survival activities in NIH 3T3 fibroblasts. Mol Cell Biol 17, 4442–4453 (1997).

42 Pao-Chun, L., Chan, P. M., Chan, W. & Manser, E. Cytoplasmic ACK1 interaction with multiple receptor tyrosine kinases is mediated by Grb2: an analysis of ACK1 effects on Axl signaling. J Biol Chem 284, 34954–34963, doi:10.1074/jbc.M109.072660 (2009).

43 Paolino, M. et al. The E3 ligase Cbl-b and TAM receptors regulate cancer metastasis via natural killer cells. Nature 507, 508–512, doi:10.1038/nature12998 (2014).

44 Holland, S. J. et al. R428, a selective small molecule inhibitor of Axl kinase, blocks tumor spread and prolongs survival in models of metastatic breast cancer. Cancer research 70, 1544–1554, doi:10.1158/0008-5472.CAN-09-2997 (2010).

45 Lew, E. D. et al. Differential TAM receptor-ligand-phospholipid interactions delimit differential TAM bioactivities. eLife 3, doi:10.7554/eLife.03385 (2014).

46 Schindeler, A., McDonald, M. M., Bokko, P. & Little, D. G. Bone remodeling during fracture repair: The cellular picture. Semin Cell Dev Biol 19, 459–466, doi:10.1016/j.semcdb.2008.07.004 (2008).

47 Xie, X. et al. Fracture risks in patients with atrial fibrillation treated with different oral anticoagulants: a meta-analysis and systematic review. Age Ageing 51, doi:10.1093/ageing/afab264 (2022).

48 Shiozawa, Y. et al. GAS6/AXL axis regulates prostate cancer invasion, proliferation, and survival in the bone marrow niche. Neoplasia 12, 116–127, doi:10.1593/neo.91384 (2010).

49 Engelmann, J. et al. Regulation of bone homeostasis by MERTK and TYRO3. Nat Commun 13, 7689, doi:10.1038/s41467-022-33938-x (2022).

50 Ryu, K. Y. et al. Mer tyrosine kinase regulates bone metabolism, and its deficiency partially ameliorates periodontitis-and ovariectomy-induced bone loss in mice. JBMR Plus 8, ziad014, doi:10.1093/jbmrpl/ziad014 (2024).

51 Maillard, C., Berruyer, M., Serre, C. M., Dechavanne, M. & Delmas, P. D. Protein-S, a vitamin K-dependent protein, is a bone matrix component synthesized and secreted by osteoblasts. Endocrinology 130, 1599–1604, doi:10.1210/endo.130.3.1531628 (1992).

52 Lemke, G. Phosphatidylserine Is the Signal for TAM Receptors and Their Ligands. Trends in biochemical sciences 42, 738–748, doi:10.1016/j.tibs.2017.06.004 (2017).

53 Krishnacoumar, B. et al. Caspase-8 promotes scramblase-mediated phosphatidylserine exposure and fusion of osteoclast precursors. Bone Res 12, 40, doi:10.1038/s41413-024-00338-4 (2024).

54 Verma, S. K. et al. Cell-surface phosphatidylserine regulates osteoclast precursor fusion. J Biol Chem 293, 254–270, doi:10.1074/jbc.M117.809681 (2018).

55 Whitlock, J. M. et al. Cell surface-bound La protein regulates the cell fusion stage of osteoclastogenesis. Nat Commun 14, 616, doi:10.1038/s41467-023-36168-x (2023).

56 Nakamura, T. et al. Estrogen prevents bone loss via estrogen receptor alpha and induction of Fas ligand in osteoclasts. Cell 130, 811–823 (2007).

57 Ferron, M., Wei, J., Yoshizawa, T., Ducy, P. & Karsenty, G. An ELISA-based method to quantify osteocalcin carboxylation in mice. Biochem Biophys Res Commun 397, 691–696 (2010).

58 Chappard, D., Palle, S., Alexandre, C., Vico, L. & Riffat, G. Bone embedding in pure methyl methacrylate at low temperature preserves enzyme activities. Acta Histochem 81, 183–190 (1987).

59 Bouxsein, M. L. et al. Guidelines for assessment of bone microstructure in rodents using micro-computed tomography. J Bone Miner Res 25, 1468–1486, doi:10.1002/jbmr.141 (2010).

60 Ferron, M. et al. Inositol polyphosphate 4-phosphatase B as a regulator of bone mass in mice and humans. Cell Metab 14, 466–477 (2011).

61 Chomczynski, P. & Sacchi, N. The single-step method of RNA isolation by acid guanidinium thiocyanate-phenol-chloroform extraction: twenty-something years on. Nature protocols 1, 581–585, doi:10.1038/nprot.2006.83 (2006).

62 Lacombe, J. et al. Vitamin K-dependent carboxylation regulates Ca(2+) flux and adaptation to metabolic stress in beta cells. Cell Rep 42, 112500, doi:10.1016/j.celrep.2023.112500 (2023).

