## Supplementary Figures 1-4 and supplementary tables 1-2 for "Vitamin K-dependent carboxylation in osteoblasts regulates bone resorption through GAS6 in male mice"

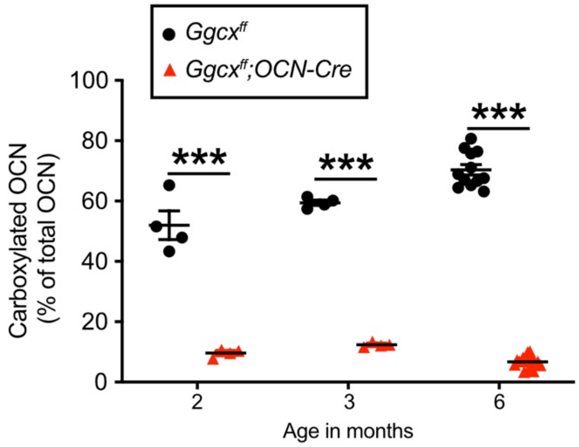


**Supplementary Figure 1. *Ggcx* inactivation in osteoblasts markedly reduced osteocalcin carboxylation level.** Serum level of carboxylated osteocalcin (OCN) expressed as a percentage of total osteocalcin in *Ggcx$^{ff}$* and *Ggcx$^{ff}$;OCN-Cre* male mice at the indicated ages (n=4-17). ***$p < 0.001$.

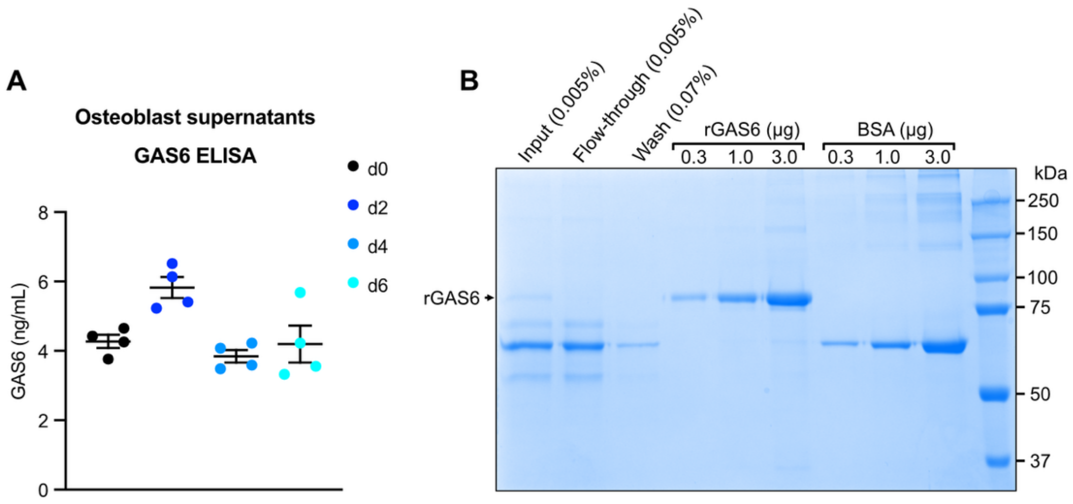


**Supplementary Figure 2. Analysis of GAS6 in osteoblast culture and purification of recombinant carboxylated GAS6. (A)** GAS6 concentration was quantified using a specific ELISA in the supernatant of osteoblasts cultured in pro-osteoclastogenic conditions (i.e., VitD$_3$ and PGE$_2$) for the indicated times (d0-d6: day 0 to 6; n=4). **(B)** Coomassie stained SDS-PAGE gel representing the different steps of the purification. Recombinant 6×HIS tagged GAS6 (rGAS6) was produced from HEK293 cells in the presence of vitamin K$_1$ to ensure maximal carboxylation. The cell supernatant (Input) was collected and GAS6 purified by nickel affinity chromatography. Following binding, GAS6 was depleted from the media (Flow-through). After extensive washes (Wash), rGAS6 was eluted using an imidazole-containing buffer, and the purified protein was dialyzed against PBS. The concentration of the purified rGAS6 was next determined using a BSA standard (BSA).

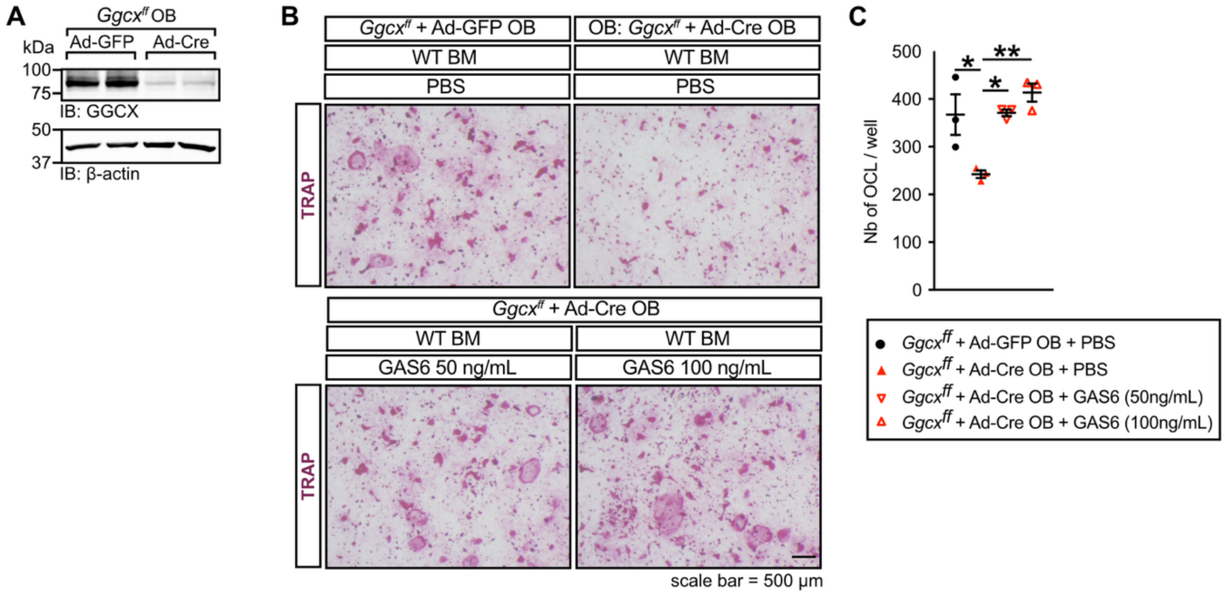


**Supplementary Figure 3. Carboxylated GAS6 rescues ex vivo osteoclastogenesis in *Ggcx* inactivated osteoblast co-cultures. (A)** Western blot analysis of extracts from *Ggcx$^{ff}$* osteoblasts transduced with Ad-GFP or Ad-Cre. **(B)** Representative TRAP staining of *Ggcx$^{ff}$* osteoblasts (OB) transduced with Ad-GFP (control) or Ad-Cre (knockout), and co-cultured with WT bone marrow cells (BM) for 7 days in the presence of prostaglandin $E_2$ ($PGE_2$; $10^{-6}$ M) and 1,25 vitamin $D_3$ ($VitD_3$; $10^{-8}$ M), with or without recombinant carboxylated GAS6 at 50ng/mL or 100ng/mL. **(C)** Quantification of the number of TRAP+ osteoclasts per well (Nb of OCL/well) (n=3). One-way ANOVA with Bonferroni's posttests was used in (C). **$p < 0.01$, *$p < 0.05$.

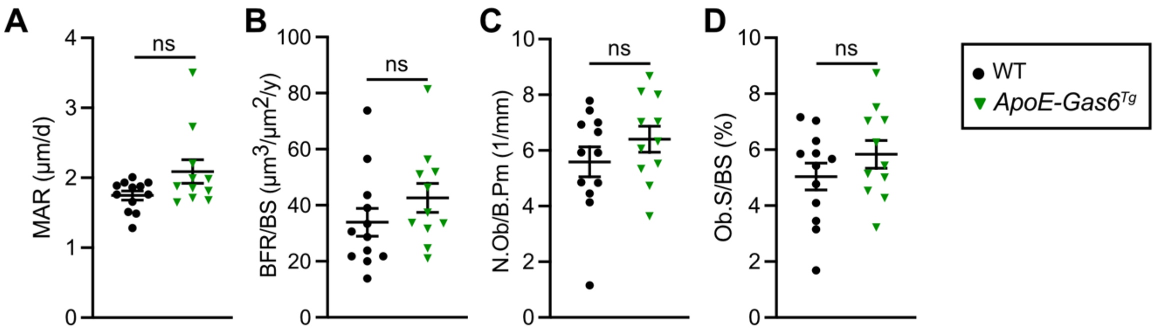


**Supplementary Figure 4. Increased circulating GAS6 level does not impact bone formation parameters.** Bone histomorphometry analysis of lumbar vertebrae in six-month-old *WT* (non-transgenic littermates) and *ApoE-Gas6*$^{Tg}$ male mice (n=11-12). **(A)** Mineral apposition rate (MAR). **(B)** Bone formation rate over bone surface (BFR/BS). **(C)** Number of osteoblasts per bone perimeter (N.Ob/B.Pm). **(D)** Osteoblast surface over bone surface (Ob.S/BS). ns: non-significant.

**Supplemental Table 1. List of oligonucleotides used in this study.**

| Gene | Forward Sequence 5'-3' | Reverse Sequence 5'-3' | Purpose |
|---|---|---|---|
| *Ggcx* | TTGACCCTCGTGTGGACATC | AATCTGCAATGAAGACCACC | qPCR |
| *Vkorc1* | ATTACCGCGCGCTCTGCGA | AAGAACAGGATCCAGGCCAG | qPCR |
| *Hprt* | TCAGTCAACGGGGGACATAAA | GGGGCTGTACTGCTTAACCAG | qPCR |
| *Rankl* | AAGATGCGACGTACTTTGGG | CGTGGGCCATGTCTCTTAGT | qPCR |
| *Opg* | GAAAGACCTGCAAATCGAGC | TTGTGAAGCTGTGCAGGAAC | qPCR |
| *Csf1* | GCAAGGGACTCACTAGCCTG | TATGCCTTTACGGGAAGTCG | qPCR |
| *Bglap* | CAGACAAGTCCCACACAGCA | CTTGGCATCTGTGAGGTCAG | qPCR |
| *Mgp* | GGCGAGCTAAAGCCCAAAAG | GTAGTCATCGCAGGCCTCTC | qPCR |
| *Ucma* | CAAGCGGTCTCCTAAGTCCC | TGGCGGTTGTAGAGGTAGGA | qPCR |
| *Postn* | AAGGCGAAACGGTGACAGAA | TCGTTCTTCCCGAGTCTGTT | qPCR |
| *F2* | TGGAAGGTCGCTGTGCTATG | CAGAGCGAGGAGTCATCACC | qPCR |
| *F7* | GAGGACTACACGCTACAGCC | CGGTCACTATCCATCTGGCG | qPCR |
| *F9* | GCAAAACCGGGTCAAATCCC | AGACAGTGGGCAGCAGTTAC | qPCR |
| *F10* | CACTGCCGTCCTTGACCAC | TTGGCACGTTCCCGGTTAAT | qPCR |
| *ProC* | GCGCTACCTGGACGAAATTG | GAACACTGGGTCAGGATGGG | qPCR |
| *ProS1* | GTGAGGGTATCCCAGTGTGC | CATCACGAAGCGCAATCAGG | qPCR |
| *ProZ* | CCCTGACTTCCGAACACATCA | CGACTCCTCGTCATAACGCAT | qPCR |
| *Gas6* | ATGAAGATCGCGGTAGCTGG | CCAACTCCTCATGCACCCAT | qPCR |
| *Prrg1* | CCAGTCACTTCCTCTGTTGGTTT | ACTAAAACGCTACCCAAGAGCC | qPCR |
| *Prrg2* | AGGCGTTTTCCTCGTGCTAA | AGGATCCCAGAGGTCAGTCC | qPCR |
| *Prrg3* | GCTCTGTGAGGGGTCTCGAA | AAGAAGCATCATGGCTGTATTTCTA | qPCR |
| *Prrg4* | CCGCCTCCTGAACAATAGGT | ATGGCCGCCTTTTACACTTG | qPCR |
| *Actb* | GACCTCTATGCCAACACAGT | AGTACTTGCGCTCAGGAGGA | qPCR |
| *MerTK* | GGTTCTGGCCCCACTGCTAC | CAGAGAATGGCCTGTGGTTGA | qPCR |
| *Axl* | CCAGTCACAGGACACAGCTC | ATACCCACCCCATCGTCTGA | qPCR |
| *Tyro3* | ACGATCTCCAGCTACAACGC | TTGTCTGAAAGGGCACCCAG | qPCR |
| *Acp5* | AGTCCTGCTTGTCCGCTAAC | CCTAAAAGGGGTGAGCCTGG | qPCR |
| *Clcn7* | TCTCGCTTGAGTGATGTTGACC | GACTGGCTGTGGGAAAGGAA | qPCR |
| *Ctsk* | CGTGCAGCAGAACGGAGGCA | GTCCTACCCGCGCCACTGCT | qPCR |
| *Dc-stamp* | TTGCCGCTGTGGACTATCTG | GAATGCAGCTCGGTTCAAAC | qPCR |
| *Ggcx* $^{flox/flox}$ | TCATTGAGTCCTTCCCGAAC | TCCAAGTGCGTCTTTAACTCC | Genotyping |
| Oc-Cre | CAAATAGCCCTGGCAGATTC | ACGCCTGGCGATCCCTGAACAT | Genotyping |
| *Bglap +/-* | TGGAGTGGTCTCTATGACCT | TTCCTTGACCCTGGAAGGTG | Genotyping |
| | TTGTGCTGGGGTGGTTTCTG | AGCCTTCCCCAACCCCTATT | |
| *Ctsk-Cre* | GCGGTCTGGCAGTAAAAACTATC | GTGAAACAGCATTGCTGTCACTT | Genotyping |
| *tdTomato* $^{flox/flox}$ | AAGGGAGCTGCAGTGGAGTA | CCGAAAATCTGTGGGAAGTC | Genotyping |
| | GGCATTAAAGCAGCGTATCC | CTGTTCCTGTACGGCATGG | |
| *ApoE-Gas6* | AAGGCTAACCTGGGGTGAGG | AAGTTCTGAACACATTTGGCGA | Genotyping |

**Supplemental Table 2. List of antibodies used in this study.**

| Antibody | Source | Catalog # | Application | Dilution |
|---|---|---|---|---|
| Rabbit anti-GGCX | ProteinTech | 16209–1-AP | WB | 1/1000 |
| Rabbit anti-VKORC1 | Ferron et al. 2015 [33] | Custom made | WB | 1/1000 |
| Mouse anti-β-Actin | MilliporeSigma | A1978 | WB | 1/2000 |
| Rabbit anti-phospho-AXL (Y702) | Schott et al. 2024 [39] | Custom made | WB | 1/1000 |
| Goat anti-Axl Antibody (C-20) | SantaCruz | sc-1096 | WB | 1/500 |
| Rabbit anti-phospho-AKT (Ser473) | Cell Signaling | 9271 | WB | 1/1000 |
| Rabbit anti-AKT (pan) (C67E7) | Cell Signaling | 4691 | WB | 1/1000 |
| Mouse anti-MERTK | Cell Signaling | 38102 | WB | 1/1000 |
